# Whole-cell scale dynamic organization of lysosomes revealed by spatial statistical analysis

**DOI:** 10.1101/219790

**Authors:** Qinle Ba, Guruprasad Raghavan, Kirill Kiselyov, Ge Yang

## Abstract

In eukaryotic cells, lysosomes are distributed in the cytoplasm as individual membrane-bound compartments to degrade macromolecules and to control cellular metabolism. A fundamental yet unanswered question is whether and, if so, how individual lysosomes are spatially organized so that their functions can be coordinated and integrated to meet changing needs of cells. To address this question, we analyze their collective behavior in cultured cells using spatial statistical techniques. We find that in single cells, lysosomes maintain nonrandom, stable, yet distinct spatial distributions, which are mediated by the coordinated effects of the cytoskeleton and lysosomal biogenesis on different lysosomal subpopulations. Furthermore, we find that throughout the intracellular space, lysosomes form dynamic clusters that substantially increase their interactions with endosomes. Together, our findings reveal the spatial organization of lysosomes at the whole-cell scale and provide new insights into how organelle interactions are mediated and regulated over the entire intracellular space.

**Highlights:**

- Lysosomes maintain stable yet distinct spatial distributions in single cells
- The cytoskeleton and lysosomal biogenesis mediate stable lysosomal distributions
- Lysosomes form dynamic clusters that promote their interactions with endosomes
- Two subpopulations of lysosomes jointly mediate formation of lysosomal clusters

**eTOC Blurb:** Lysosomes are spatially organized at the whole-cell scale and form dynamic clusters that promote their interactions with endosomes.

## INTRODUCTION

A basic strategy used by eukaryotic cells to organize their internal environment is to form specialized membrane-bound organelles such as lysosomes and endosomes. Although this strategy provides important structural and functional benefits, specialized functions of the organelles must be coordinated and integrated for cell physiology (Gottschling and Nyström, 2017). Recent studies have shown that different organelles interact directly and extensively through mechanisms such as membrane contact (Prinz, 2014, Helle et al., 2013) and membrane fusion (Martens and McMahon, 2008, McNew et al., 2013). Such interactions depend critically on the colocalization and, therefore, the spatial distributions of the organelles. Currently, however, we cannot explain how they are mediated and regulated at the systems level within the dynamic and heterogeneous intracellular space.

To address this deficiency, we focus specifically on the lysosome, an organelle that plays essential roles in important cellular functions such as degrading macromolecules (Luzio et al., 2007, Xu and Ren, 2015) and controlling cellular metabolism (Lim and Zoncu, 2016, Settembre et al., 2013). Within the intracellular space, lysosomes are distributed as individual compartments. Individual lysosomes are limited in their own capacity. The size of a lysosome is typically limited to several hundred nanometers (Yu et al., 2010, Xu and Ren, 2015, Bandyopadhyay et al., 2014). The number of lysosomes in a mammalian cell is typically limited to several hundred (Valm et al., 2017, Xu and Ren, 2015). Furthermore, lysosomes are specialized compartments and dependent on interactions with partner organelles to fulfill their functions (Luzio et al., 2007, Bonifacino and Neefjes, 2017). For example, they depend on fusion with endosomes and autophagosomes to receive and degrade materials from the endocytic and autophagic pathways, respectively (Luzio et al., 2007, Eskelinen and Saftig, 2009). Given the functional limitations of individual lysosomes, a fundamental yet unanswered question is whether and, if so, how individual lysosomes are organized in the intracellular space so that their functions can be coordinated and integrated to meet changing needs of cells. Answering this question is key to elucidating how lysosomes function in single cells at the systems level.

Recent studies on positioning of lysosomes (Bonifacino and Neefjes, 2017, Pu et al., 2016) have started to reveal their spatial organization within the intracellular environment. Positioning of individual lysosomes is mediated by mechanisms including their active transport along microtubules as well as their interactions with the actin cytoskeleton and partner organelles, especially the endoplasmic reticulum (Pu et al., 2016, Bonifacino and Neefjes, 2017). Under normal conditions, lysosomes in non-polarized mammalian cells often cluster in a perinuclear region surrounding the microtubule-organizing center (MTOC), forming what is called the perinuclear cloud (Pu et al., 2016, Jongsma et al., 2016, Korolchuk et al., 2011). But they also spread into peripheral regions of cells, with some approaching the plasma membrane. This spatial pattern provides direct evidence for spatial organization of individual lysosomes, and a wide variety of perturbations can change this pattern (Pu et al., 2016). For example, nutrient deprivation substantially increases the fraction of lysosomes clustering in the perinuclear region and decreases the fraction of lysosomes spreading into peripheral regions. Nutrient recovery reverses these changes and restores the usual pattern of lysosomal distribution (Korolchuk et al., 2011, Li et al., 2016). The relocation of lysosomes under these conditions is mediated by motor-mediated active transport along microtubules, and the underlying molecular machineries and mechanisms have started to be elucidated (Korolchuk et al., 2011, Li et al., 2016, Pu et al., 2015). Functions of lysosomal positioning in mediating cellular nutrient response (Korolchuk et al., 2011) and regulating lysosomal degradative capacity (Johnson et al., 2016) have also started to be elucidated. Despite these advances, related studies have a basic limitation in elucidating the spatial organization of lysosomes in that they lack quantitative and comprehensive characterization and analysis of the collective behavior of lysosomes, especially at the whole-cell scale.

That subcellular structures such as organelles and proteins exhibit defined spatial patterns has been noted in many studies (Valm et al., 2017, Boland and Murphy, 2001, Glory and Murphy, 2007). These patterns reflect the spatial organization of the subcellular structures and have been characterized and analyzed using pattern recognition and machine learning techniques (Boland and Murphy, 2001, Johnson et al., 2015, Li et al., 2012, Naik et al., 2016). However, the specific modes, molecular mechanisms, and cellular functions of these patterns remain poorly understood.

In this study, we probe the spatial organization of lysosomes in cultured COS-7 or BS-C-1 cells. Our overall strategy is to study collective behavior of lysosomes, especially their spatial distribution at the whole-cell scale, using spatial statistical analysis techniques. Specifically, we treat the spatial distribution of lysosomes mathematically as a spatial point process and analyze it using related statistical techniques (Diggle, 2014, Illian et al., 2008, Baddeley et al., 2016). We find that lysosomes maintain nonrandom, stable, yet distinct spatial distributions in single cells, confirming that they are spatially organized at the whole-cell scale. We then probe the molecular mechanisms underlying the stable distributions. We find that they are maintained by the coordinated effects of several mechanisms, including microtubule-based active transport, interaction with the actin cytoskeleton, and lysosomal biogenesis, on different lysosomal subpopulations. Lastly, we investigate the relation between the interactions of lysosomes with late endosomes and their spatial distributions. We find that throughout the intracellular space, lysosomes form dynamic clusters that locally and substantially increase their interactions with endosomes without increasing the numbers of these organelles globally. Furthermore, we find that formation of the clusters is mediated primarily by lysosomes undergoing motor-mediated directed movement together with lysosomes undergoing constrained diffusion. Together, our findings identify specific modes, molecular mechanisms, and cellular functions of the spatial organization of lysosomes at the whole-cell scale and provide new insights into how cells organize their organelles and mediate their interactions.

## RESULTS

### Spatial distribution of lysosomes at the whole-cell scale is nonrandom and stable

To investigate whether and, if so, how lysosomes are spatially organized, we analyzed their spatial distribution at the whole-cell scale in live BS-C-1 cells. We labelled them using dextran Alexa-488 and collected time-lapse movies at four frames per second for one minute (Fig. 1A). Within each cell we analyzed, we observed extensive movement of lysosomes throughout the intracellular space, with many traversing long distances over the duration of imaging (Fig. 1B; Supplementary Movie S1). For each cell we analyzed, we located its individual lysosomes in each frame as single particles using image analysis software (Materials & Methods). We then considered their spatial distribution mathematically as a spatial point process and analyzed it using related spatial statistical techniques (Illian et al., 2008, Diggle, 2014, Baddeley et al., 2016). Specifically, we checked whether the lysosomes were randomly distributed by performing complete spatial randomness (CSR) tests (Materials & Methods) within the cell boundary at different time points throughout the movie. We found that the spatial distribution of lysosomes differed substantially from a random distribution at all the time points we analyzed (Fig. 1C-E). This shows that lysosomes are spatially organized at the whole-cell scale.

**Figure 1.**
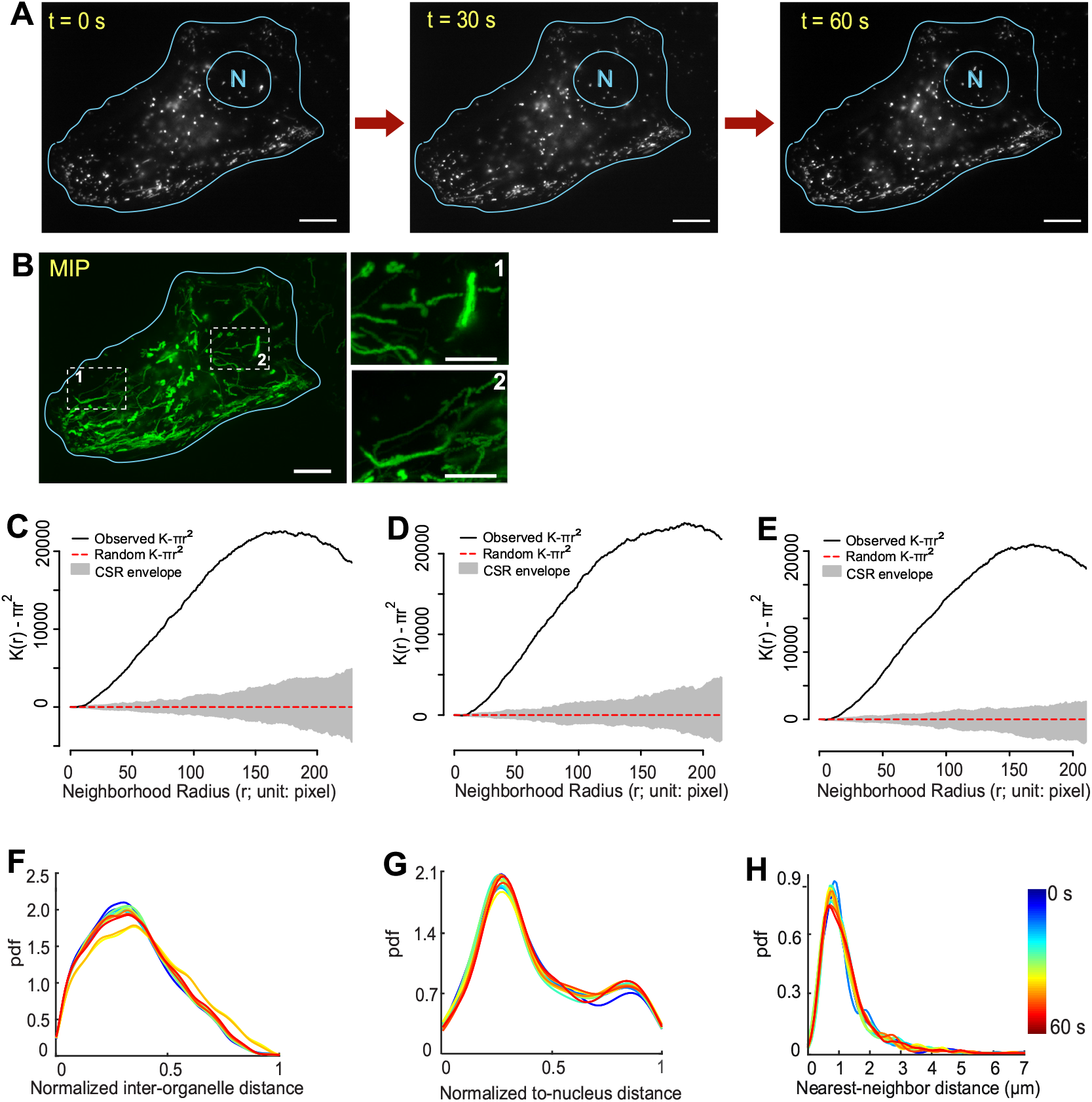
Lysosomes maintain a nonrandom and stable spatial distribution in a single cell. (A) Lysosomes in a BS-C-1 cell at three selected time points. N: nucleus. Scale bars: 10 µm. (B) Maximum intensity projection (MIP), shown in green, of lysosomal movement over one minute, imaged at four frames per second. Each trace corresponds to the trajectory of a lysosome. Scale bar: 10 µm. Scale bars in insets: 5 µm. (C-E) Complete spatial randomness (CSR) test of whole-cell scale lysosomal distribution at the three time points in (A), respectively. (C):0 s; (D):30 s; (E):60 s. Solid black line: adjusted Ripley’s *K*-function of lysosomes within the cell. Dotted red line: adjusted Ripley’s *K*-function of a random distribution within the same cell boundary. Gray area: uncertainty envelope for the random distribution. The extent of separation between the solid black line and the CSR envelope indicates how close the spatial distribution of lysosomes is to a random distribution. (F-H) Three distance distributions of lysosomes, color-coded based on time and plotted every 5 seconds over 60 seconds. Their temporal variations were quantified using Sorensen dissimilarity scores. Because the distributions at 0, 5, 10, 15,…, 60 seconds were selected, 13 distributions were compared pairwise, hence 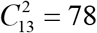 pairs. pdf: probability density function. (F-H) Temporal variations (mean ± STD; n = 78). (F) Normalized inter-organelle distance distribution: 3.29% ± 2.98%. (G) Normalized to-nucleus distance distribution: 2.65% ± 0.92%. (H) Nearest-neighbor distance distribution: 5.80% ± 1.48%.

To quantitatively characterize the spatial distribution of lysosomes at the whole-cell scale, we used three statistical distance distributions (Fig. S1A-C; Materials & Methods). We calculated these distributions every five seconds in each cell so that we could examine their variations over time (Fig. 1F-H). First, we quantified the positioning of all the lysosomes relative to each other in each cell using the distribution of their normalized pairwise distances (Fig. S1A), which we refer to as the normalized inter-organelle distance distribution (Diggle, 2014). For the cell shown in Fig. 1A-B, this distribution remained generally stable over the duration of imaging, whose profile spread broadly but showed a peak and thus a characteristic spacing at ∽0.35 (Fig. 1F). The distributions in other cells we analyzed showed similar stability and comparable but different profiles (Fig. 2A blue lines; Fig. S1D-E). Second, we quantified the positioning of all the lysosomes relative to the nucleus using the distribution of their normalized shortest distances to the nucleus (Fig. S1B), which we refer to as the normalized to-nucleus distance distribution (Fig. 1G). For the cell shown in Fig. 1A-B, this distribution also remained generally stable, but its profile showed two peaks and therefore two characteristic distances, a primary one at ∽0.3 and a secondary one at ∽0.9. The distributions in other cells showed similar stability but remarkably diverse profiles (Fig. 2B blue lines; Fig. S1D-E). Third, we quantified the level of crowding of the lysosomes in each cell, i.e. how closely they are spaced, using the distribution of their nearest-neighbor distances (Fig. S1C), which we refer to as the nearest-neighbor distance distribution (Fig. 1H) (Illian et al., 2008, Diggle, 2014, Baddeley et al., 2016). For the cell shown in Fig. 1A-B, the distribution also remained stable and showed a characteristic peak distance at ∽1 µm (Fig. 1H). The distributions in other cells showed similar stability and similar profiles (Fig. 2C blue lines; Fig. S1D-E). Together, the three distance distributions provide a comprehensive characterization of the spatial distribution of lysosomes at the whole-cell scale.

**Figure 2.**
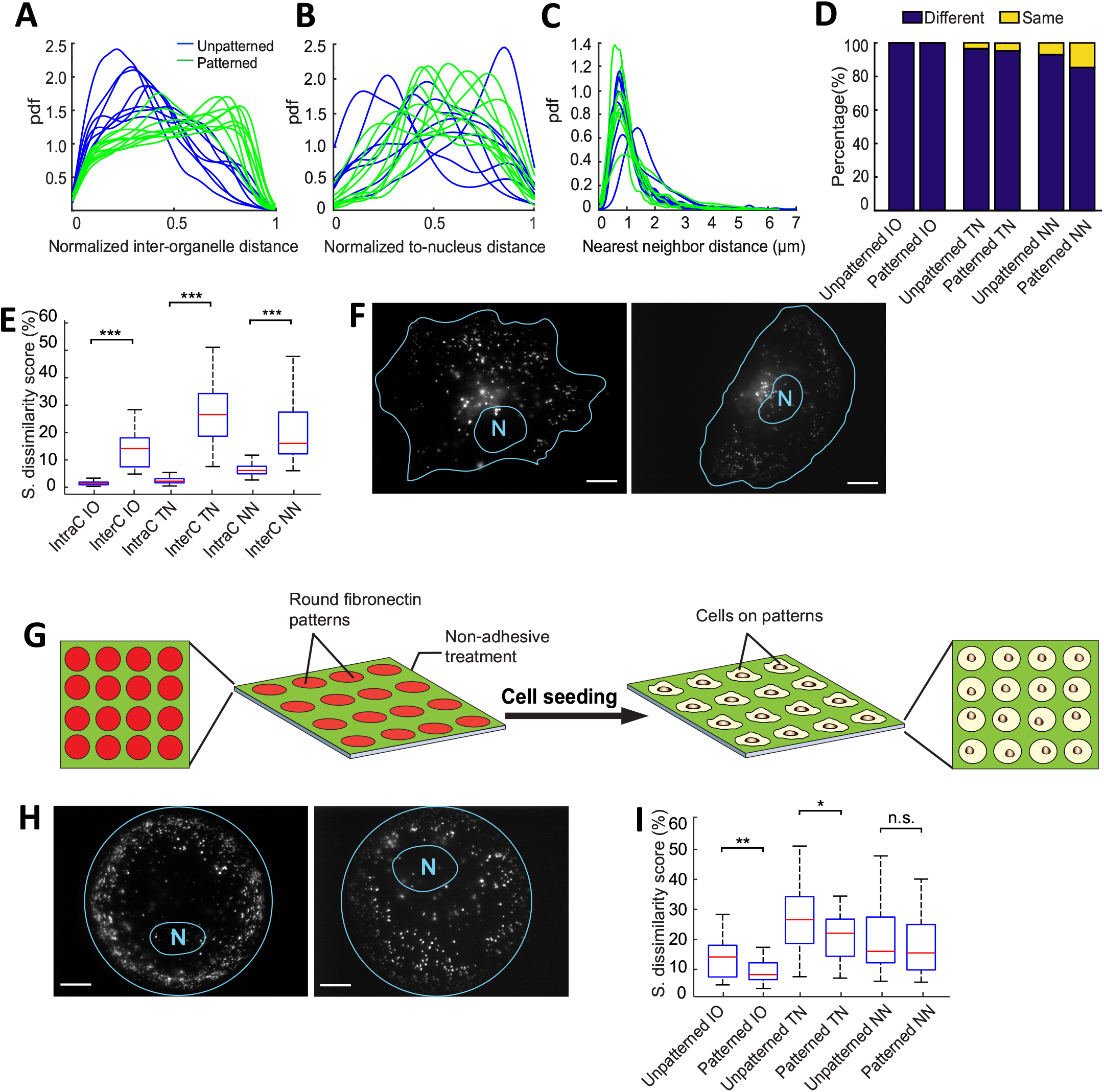
Distinct lysosomal distributions in singles cells are not merely a secondary effect of their distinct shapes. (A-C) Comparison of normalized inter-organelle distance distributions (A), normalized to-nucleus distance distributions (B), and nearest-neighbor distance distributions (C) in non-patterned cells (blue lines; n = 8) and patterned cells (green lines; n = 10). (D) Results of pairwise comparison of the three distance distributions among non-patterned (total = 28 comparisons; left columns) and patterned cells (total = 45 comparisons; right columns) using two-sample Kolmogorov-Smirnov tests. Cutoff p-value for statistical significance: 0.05. IO: normalized inter-organelle distance distribution; TN: normalized to-nucleus distance distribution. NN: nearest-neighbor distance distribution. Percentage of comparison showing significant differences: IO: 100% (non-patterned) and 100% (patterned); TN: 96.4% (non-patterned) and 95.6% (patterned); NN: 92.9% (non-patterned) and 84.4% (patterned). (E) Comparison of intracellular (intraC) variations of the distance distributions within single cells versus intercellular (interC) variations of the distributions among different cells using Wilcoxon rank sum tests. All variations represented in Sorensen dissimilarity scores. IO: intracellular: 1.64% ± 1.38% (mean ± STD; n = 624 scores; data pooled from 8 cells with 78 scores per cell), intercellular: 14.42% ± 7.27% (mean ± STD; n = 28 scores from 8 cells), p-value: 1.3×10^-18^; TN: intracellular: 2.67% ± 1.80% (n = 624), intercellular: 27.89% ± 12.83% (n = 28), p-value: 4.5×10^-19^; NN: intracellular: 6.46% ± 2.14% (n = 624), intercellular: 21.53% ± 13.42% (n = 28), p-value: 1.2×10^-15^. Notation for p values: *, p < 0.05; **, p < 0.01; ***, p < 0.001. (F) Examples of unpatterned cells with different shapes. Scale bars: 10 µm. (G) A cartoon illustrating the process of patterning shapes of cells by growing them on patterned fibronectin substrates. (H) Examples of patterned cells. Scale bars: 10 µm. (I) Comparison of intercellular variations of non-patterned cells versus patterned cells using Wilcoxon rank sum tests. IO: non-patterned: 14.42% ± 7.27% (mean ± STD; n = 28 scores from 8 cells); patterned: 9.42% ± 3.99% (mean ± STD; n = 45 scores from 10 cells), p-value: 0.0037. TN: non-patterned: 27.89% ± 12.83% (n = 28); patterned: 20.65% ± 7.63% (n = 45), p-value: 0.016. NN: non-patterned: 21.53% ± 13.42% (n = 28); patterned: 18.94% ± 10.50% (n = 45), p-value: 0.59.

We have noted that the three distance distributions remained generally stable over the duration of imaging in each cell we analyzed. To quantitatively characterize the stability of each distribution, we quantified its variations over time using the Sorensen dissimilarity score between its profiles at any two different selected time points (Materials & Methods). This score is also referred to as the Sorensen distance between two probability density distributions (Cha, 2007). For the cell shown in Fig. 1A-B, the average dissimilarity score was 3.29% for the normalized inter-organelle distance distribution, 2.65% for the normalized to-nucleus distance distribution, and 5.80% for the nearest-neighbor distance distribution. Examination of these distributions in different cells (Fig. S1D-E) and over different durations of imaging (Fig. S1F-G) found similarly low levels of variations. Taken together, our data show that, despite the extensive long-distance movement of lysosomes over the entire intracellular space, their spatial distribution at the whole-cell scale is nonrandom and stable, indicating homeostasis in their positioning relative to each other and to the nucleus in single cells. This provides further evidence that lysosomes are spatially organized.

### Distinct lysosomal spatial distributions in single cells are not merely a secondary effect of distinct cell shapes

We next compared the spatial distributions of lysosomes in different cells. Because the distribution within each cell remained stable over time, we selected the first frames from time-lapse movies of different cells and compared the three lysosomal distance distributions among all cells in a pairwise fashion (Fig. 2A-C; blue lines). We found that the normalized inter-organelle distributions and the normalized to-nucleus distance distributions differed significantly in all or nearly all pairwise comparisons (Fig. 2D, left columns). For the nearest-neighbor distance distributions, only ∽7% of the pairwise comparisons showed no significant difference (Fig. 2D, left column). Taken together, our data show that lysosomes maintain distinct spatial distributions in single cells.

To investigate what causes the variations among single cells in their lysosomal distributions, we examined two sources. First, temporal variations of the three distance distributions within single cells, which we refer to as intracellular variations, surely contribute to the variations among different cells, which we refer to as intercellular variations. Overall, however, we found that the contribution was very small because the average level of intracellular variations was significantly lower than the average level of intercellular variations (Fig. 2E). Second, different cells often exhibit distinct shapes (Fig. 2F). To check whether variations among different cells in their lysosomal distributions are merely a secondary effect of variations in their shapes, we grew cells into approximately the same size and circular shape on patterned protein substrates (Fig. 2G-H; Materials & Methods; Supplementary Movie S2). We calculated the three distance distributions for the patterned cells (Fig. 2A-C; green lines) and then checked the intercellular variations. All or most of the pairwise comparisons of these distributions showed significant difference (Fig. 2D, right columns). For the normalized inter-organelle distance distribution and the normalized to-nucleus distance distribution, the levels of intercellular variations among patterned cells remained substantial, thought significantly reduced from the levels of intercellular variations among unpatterned cells (Fig. 2I). For the nearest-neighbor distance distribution, there was no significant difference between unpatterned and patterned cells in terms of intercellular variations (Fig. 2I). Taken together, our results show that, although variations in cell shapes are a contributing factor to intercellular variations in spatial distributions of lysosomes, distinct lysosomal spatial distributions in unpatterned cells are not just a secondary effect of the distinct cell shapes. Instead, our results suggest that the stable yet distinct spatial distributions of lysosomes in single cells are mediated by intrinsic intracellular mechanisms. This further indicates that lysosomes are spatially organized.

### Lysosomes in a single cell form different subpopulations

Our study thus far has focused on the spatial distribution of the entire population of lysosomes in single cells. However, it is clear even from a simple visual inspection that the population is heterogeneous in its dynamic behavior: while some lysosomes traverse long distances, others seem constrained in their movement (Fig. 3A; Supplementary Movie S1). This raises the important question of how lysosomes with different behaviors together maintain the stable spatial distribution of their whole population. To answer this question, we first examined the composition of the lysosomal population in COS-7 cells by tracking individual lysosomes as single particles and characterizing their behavior through mean square displacement (MSD) analysis (Fig. 3B-C) (Qian et al., 1991, Saxton, 1997, Metzler and Klafter, 2000). We found that on average, ∽49% of lysosomes in a single cell underwent constrained diffusion, ∽31% of lysosomes underwent directed movement, and ∽20% of lysosomes underwent free diffusion (Fig. 3C). To give a concrete example of how far these subpopulations of lysosomes travel, we calculated their mean displacement over 5 seconds and found that lysosomes undergoing constrained diffusion, free diffusion, and directed movement traversed an average of 0.24 ± 0.17µm (mean ± STD; n = 768 trajectories from 9 cells), 0.50 ± 0.34 µm (n = 512 trajectories from 9 cells), and 1.20 ± 0.84 µm (n= 356 trajectories from 9 cells), respectively. Together, these results show that most lysosomes undergo either constrained diffusion or directed movement, and only a small fraction undergoes free diffusion, consistent with the finding of previous studies that free diffusion is limited inside cells (Bandyopadhyay et al., 2014, K Luby-Phelps et al., 1988, Luby-Phelps et al., 1987).

**Figure 3.**
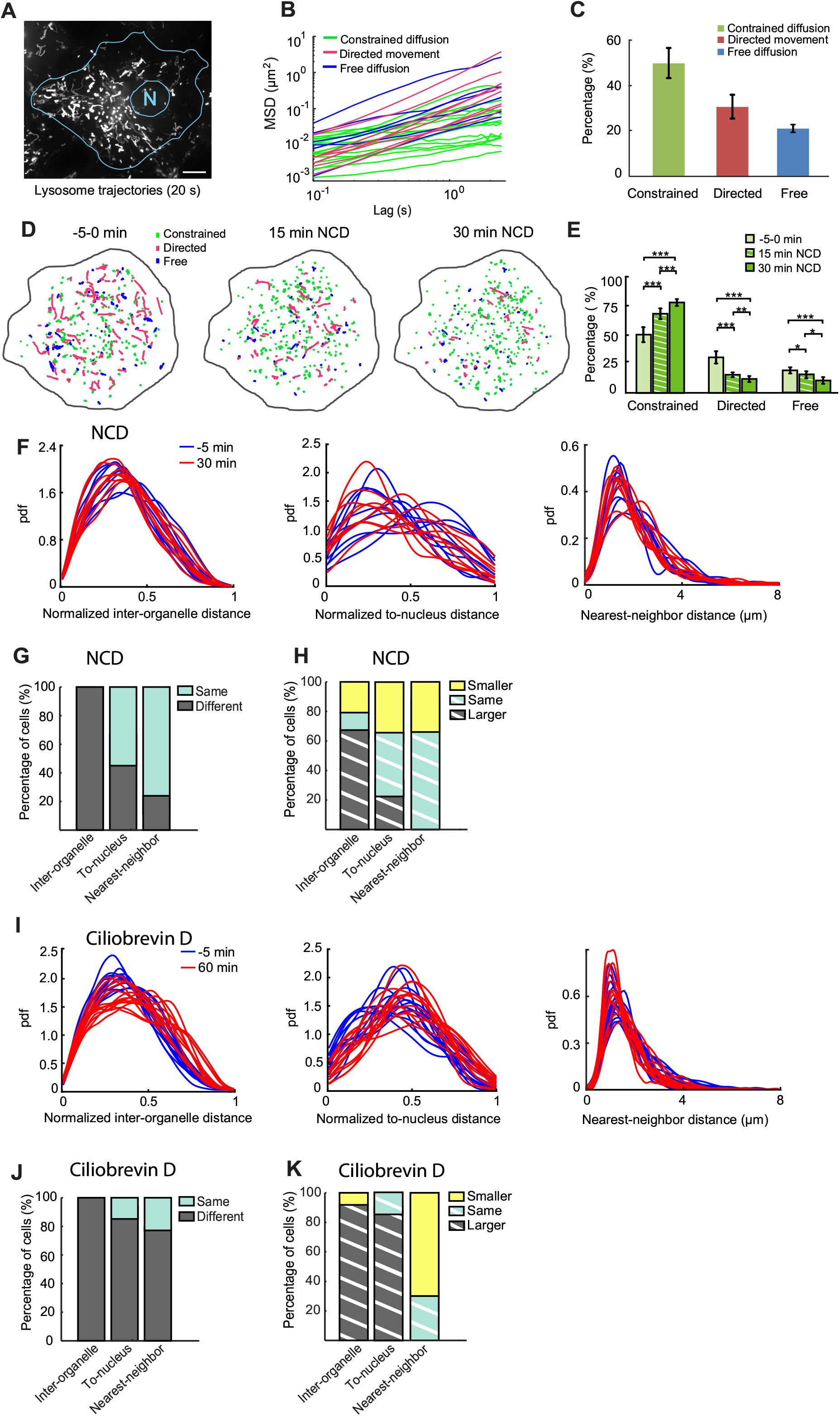
Composition of the lysosomal population and roles of microtubule-based active transport in maintaining its stable spatial distribution. (A) Maximum intensity projection of movement of mCherry-Lamp1 labeled lysosomes in a COS-7 cell from a time-lapse movie collected at 10 frames per second for 20 seconds. Scale bar: 15 µm. (B) MSD of randomly selected 10% of all trajectories within a COS-7 cell. (C) Percentage of each subpopulation (mean±STD; n = 9 cells): constrained diffusion: 49.41%±6.24%; directed movement: 30.94%±5.40%; free diffusion: 19.64%±2.19%. (D) Color-coded trajectories of lysosomes at three time points in a cell treated with 2.5 uM nocodazole (NCD). (E) Changes in lysosomal subpopulations under nocodazole treatment over time. Constrained diffusion: 49.41%±6.24% (mean±STD; n = 9 cells; 0 min), 67.74%±4.17% (n = 9 cells; 15 min), 77.20%±3.32% (n = 9 cells; 30min). Directed movement: 30.94%±5.40% (0 min), 16.77%±2.05% (15 min), 11.71%±2.48% (30min). Free diffusion: 19.65%±2.19% (0 min), 15.49%±3.32% (15 min), 11.05%±3.80% (30min). Comparison was made using two sample student-t tests on pooled data from the same cells before and after NCD treatment. p-values: constrained diffusion: 1.8×10^-4^ (0 min vs 15 min), 9.6×10^-6^ (0 min vs 30 min), 8.1×10^-4^ (15 min vs 30 min); directed movement: 2.7×10^-5^ (0 min vs 15 min), 3.2×10^-5^ (0 min vs 30 min), 2.0×10^-3^ (15 min vs 30 min); free diffusion: 2.6×10^-2^ (0 min vs 15 min), 7.3×10^-5^ (0 min vs 30 min), 1.3×10^-2^ (15 min vs 30 min) (F) The three distance distributions before and after 30 minutes of NCD treatment in 9 different cells. (G) Comparison of different distance distributions of lysosomes before versus after NCD treatment using two sample Kolmogorov-Smirnov tests. Cutoff p-value for statistical significance: 0.05. Normalized inter-organelle distance distribution: same 0%, different 100%; Normalized to-nucleus distance distribution: same 55.5%, different 44.4%; Nearest-neighbor distance distribution: same 77.8%, different 22.2%. (H) Comparison of median distances before versus after NCD treatment using Wilcoxon rank sum tests. Normalized inter-organelle distance distribution: smaller 22.2%, same 11.1%, larger 66.7%; normalized to-nucleus distance distribution: smaller 33.3%, same 44.4%, larger 22.2%; nearest-neighbor distance distribution: smaller 33.3%, same 66.7%, larger 0%. (I) The three distance distributions before and after 1 hour of ciliobrevin (80 μM) treatment in 13 different cells. (J) Comparison of three distance distributions before versus after ciliobrevin D treatment (80 μM, 1 hour) of the same cells using two sample Kolmogorov-Smirnov tests: Normalize inter-organelle distance distribution: same 0%, different 100%; normalized to-nucleus distance distribution: same 15.4%, different 84.6%; nearest-neighbor distance distribution: same 23.1%, different 76.9%. (K) Comparison of median distances before versus after ciliobrevin D treatment (80 µM, 1 hour) using Wilcoxon rank sum tests. Normalized inter-organelle distance distribution: smaller: 7.7%; same: 0%, larger 92.3%; normalized to-nucleus distance distribution: smaller 0%, same 15.4%, larger 84.6%; nearest-neighbor distance distribution: smaller 69.2%, same 30.8%, larger 0%.

### Sustained and balanced transport along microtubules is required for maintaining population composition and spatial distributions of lysosomes

After determining the composition of the lysosomal population, we investigated the relation between microtubule-based active transport and the different subpopulations of lysosomes. To this end, we depolymerized the microtubule cytoskeleton by treating COS-7 cells with 2.5 µM of nocodazole (Supplementary Movie S3). After 15 minutes of treatment, the fraction of lysosomes undergoing directed movement was significantly reduced, from 31% to 17% (Fig. 3D-E). After 30 minutes of treatment, the fraction was further reduced to ∽12% (Fig. 3D-E). In the meantime, the fraction of lysosomes undergoing constrained diffusion was significantly increased, from 49% to 77% after 30 minutes of treatment, while the fraction of lysosomes undergoing free diffusion was significantly reduced from 20% to 11% (Fig. 3D-E). These data show that inhibition of microtubule-based transport causes a significant fraction of lysosomes to switch from directed movement to constrained diffusion. It can also be inferred from these data that after their directed movement, lysosomes mostly switch to constrained diffusion rather than free diffusion. Given that in control cells the total number of lysosomes (Fig. S2) and the population composition remain stable, the number of lysosomes switched from directed movement to constrained diffusion should be balanced by approximately the same number switching vice versa. Taken together, these results show that sustained microtubule-based transport is responsible for maintaining the lysosomal subpopulation undergoing directed movement as well as its balance with the subpopulation undergoing constrained diffusion.

Because microtubule-based active transport is crucial to the positioning and relocation of lysosomes (Bonifacino and Neefjes, 2017, Pu et al., 2016), we reason that it should play an important role in maintaining the stable spatial distribution of lysosomes. To test this hypothesis, we compared the three distance distributions of the lysosomes right before and 30 minutes after the treatment (Fig. 3F). We found that although inhibition of the transport caused significant changes to the normalized inter-organelle distribution in all the cells we analyzed (Fig. 3G; Table S1), it did not consistently decrease or increase the median distance between lysosomes (Fig. 3H; Table S1). Analysis of the normalized to-nucleus distance distribution showed that it remained unchanged in ∽56% of the cells (Fig. 3H) and that inhibition of the transport did not consistently decrease or increase of its median distance either (Fig. 3H). Lastly, the nearest-neighbor distance distribution and its median distance remained unchanged in the majority of the cells, indicating that microtubule-based transport does not play a major role in maintaining the crowding of lysosomes.

We hypothesize that the lack of consistent changes in the median normalized inter-organelle distance and the median normalized to-nucleus distance after nocodazole treatment is because the centrifugal transport and the centripetal transport of the lysosomal population are balanced. Abolishment of the entire transport process by nocodazole treatment would not shift the balance consistently in either direction. To test this hypothesis, we treated the cell with dynein inhibitor ciliobrevin D (Firestone et al., 2012), which disrupts primarily the centripetal movement and shifts the balance towards the centrifugal movement. We found that inhibiting dynein-mediated transport caused significant changes to the three distance distributions in all or most of the cells analyzed (Fig. 3I-K; Table S2). It also consistently increased the median normalized to-nucleus distance and the median normalized inter-organelle distance in most of the cells and decreased the median nearest-neighbor distance in the majority of cells (Fig. 3I-K; Table S2). Taken together, our findings indicate that the balance between centrifugal and centripetal transport is required for maintaining stable spatial distributions of lysosomes. Because of this balance, lysosomes can undergo long distance transport without affecting their overall stable spatial distribution at the whole-cell scale.

### Interaction with the actin cytoskeleton is required for constraining diffusion of lysosomes and maintaining their spatial distributions

We have shown that microtubule-based transport is responsible for maintaining the lysosomal subpopulation undergoing directed movement. This raises the question of what maintains the subpopulation undergoing constrained diffusion. Interaction of lysosomes with the actin cytoskeleton plays an important role in mediating their positioning (Pu et al., 2016, Bonifacino and Neefjes, 2017). In particular, interaction with the cortical actin network has been shown to transiently constrain lysosomes near the cell periphery (Caviston et al., 2011, Encarnação et al., 2016). We hypothesize that interaction with the actin cytoskeleton is responsible for maintaining the subpopulation of lysosomes undergoing constrained diffusion.

To test this hypothesis, we depolymerized the actin cytoskeleton by treating cells with 0.8 µM of latrunculin A (latA). We observed substantial shape changes in many cells after 15-16 minutes of treatment, indicating that the depolymerization was effective (Fig. S3A). To avoid complications in result interpretation, we chose to analyze cells without visible changes in their shapes. To determine how depolymerization of the actin cytoskeleton affected the three lysosomal subpopulations, we tracked individual lysosomes and calculated their mean displacement over 5 seconds immediately before the treatment, 7-8 minutes after the treatment, and 15-16 minutes after the treatment (Fig. 4A-B; Supplementary Movie S4). We observed significant increases in mean displacements of all three subpopulations (Fig. 4B; Fig. S3B; Table S3). Despite these significant changes, the fraction of lysosomes undergoing constrained diffusion was only slightly reduced, from 42.1% to 38.9% after 15-16 minutes of treatment (Fig. 4C). Correspondingly, the fraction of lysosomes undergoing directed movement was slightly increased, from 37.5% to 39.3%, while the fraction of lysosomes undergoing free diffusion remained unchanged (Fig. 4C). We further checked and confirmed that our latA treatment effectively depolymerized the actin cytoskeleton using immunofluorescence (Fig. S3C-D). Together, our data show that interaction with the actin cytoskeleton constrains diffusion of lysosomes. However, it plays only a minor role in maintaining the population composition of lysosomes (Fig. 4C). We propose that the composition is defined primarily by the attachment and detachment of lysosomes with microtubules via molecular motors and adapters, in which interaction with the actin cytoskeleton plays a minor role.

**Figure 4.**
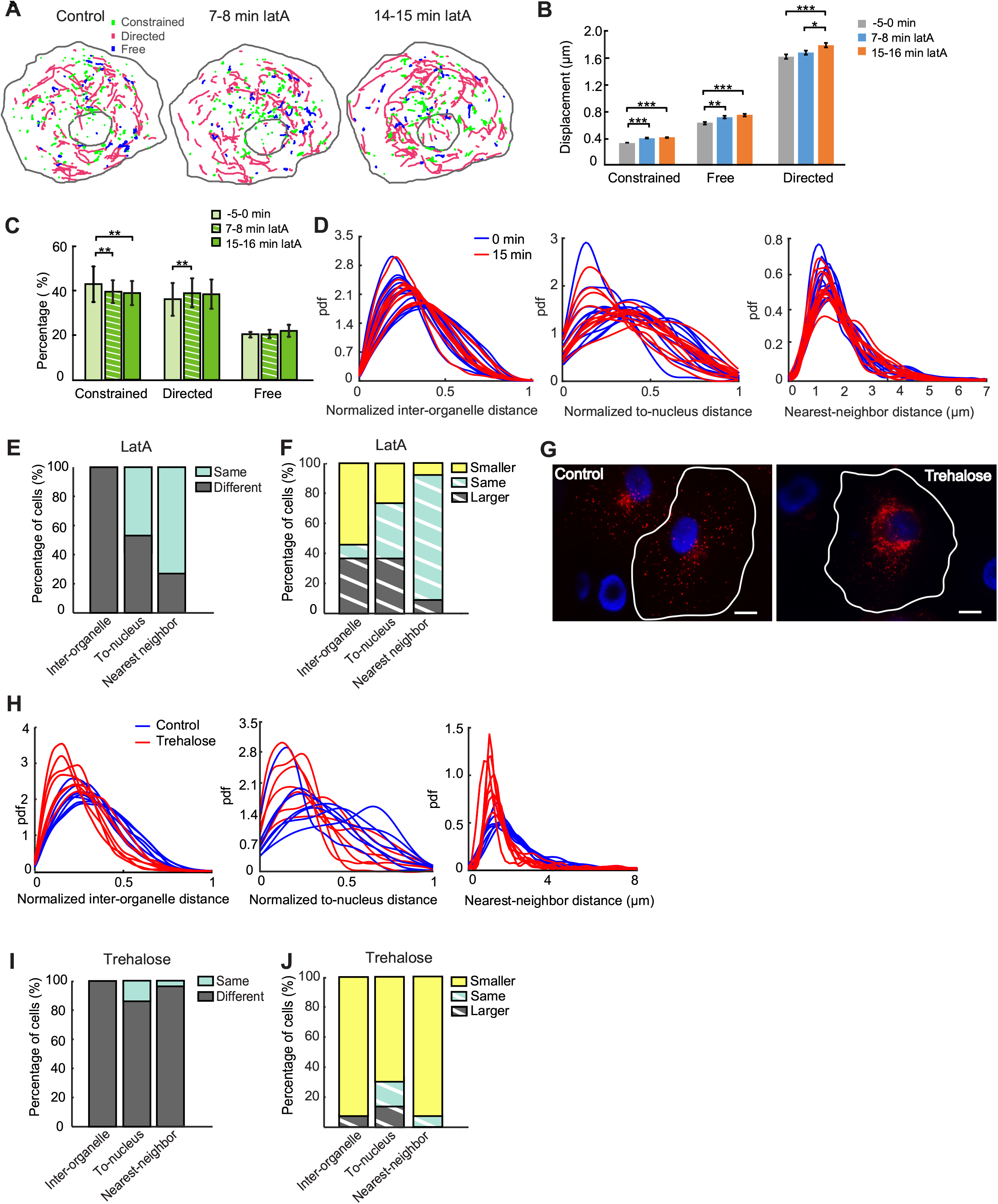
Roles of interaction with the actin cytoskeleton and lysosomal biogenesis in maintaining subpopulations and spatial distribution of lysosomes. (A) Color-coded trajectories of lysosomes in cells transfected with mCherry-Lamp1 and treated with 0.8 µM latrunculin A. Movies were collected at 10 frames per second for 20 seconds. (B) Changes in the displacement over 5 seconds of the three subpopulations under latA treatment. Comparison was made using two sample student-t tests on pooled data from the same 11 cells before and after latrunculin A treatment Constrained diffusion: 0.32±0.0058 μm (mean±SEM; 0 min, n = 2052 trajectories), 0.39±0.0076 μm (7.5 min, n = 1821 trajectories), 0.41±0.0075 μm (15 min, n = 1747 trajectories). Free diffusion: 0.57±0.019 μm (0 min, n = 688 trajectories), 0.66±0.020 μm (7.5 min, n = 633 trajectories), 0.68±0.019 μm (15 min, n = 675 trajectories). Active transport: 1.61±0.031 μm (0 min, n = 1757 trajectories), 1.67±0.031 μm (7.5 min, n = 1833 trajectories), 1.78±0.032 μm (15min, n = 1713 trajectories). p-values: constrained diffusion: 1.2×10^-14^ (0 min vs 7.5 min), 1.9×10^-22^ (0 min vs 15 min), 0.083 (7.5 min vs 15 min); free diffusion: 2.1×10^-3^ (0 min vs 7.5 min), 3.6×10^-5^ (0 min vs 15 min), 0.32 (7.5 min vs 15 min); directed movement: 0.17 (0 min vs 7.5 min), 1.6×10^-4^ (0 min vs 15 min), 0.014 (7.5 min vs 15 min). (C) Changes in the three subpopulations under latA treatment over time (mean±STD; n = 11 cells). Constrained diffusion: 42.14%±6.90% (0 min), 39.34%±5.09% (7.5 min), 38.96%±5.60% (15 min). Directed movement: 37.52%±7.02% (0 min), 40.25%±6.00% (7.5 min), 39.30%±6.20% (15min). Free diffusion: 20.34%±1.31% (0 min), 20.41%±1.68% (7.5 min), 21.74%±2.51% (15min). Comparison was made using two sample student-t tests. p-values: constrained diffusion 0.0058 (0 min vs 7.5 min), 0.0053 (0 min vs 15 min), 0.49 (7.5 min vs 15 min); directed movement 0.0043 (0 min vs 7.5 min), 0.10 (0 min vs 15 min), 0.61 (7.5 min vs 15 min); free diffusion 0.98 (0 min vs 7.5 min), 0.15 (0 min vs 17 min), 0.22 (15 min vs 15 min); (D) The three distance distributions before and 15 minutes after latA treatment in 11 cells. (E) Comparison of distributions before and after latA treatment using two sample Kolmogorov-Smirnov tests (n = 55 pairs): Inter-organelle distance distribution: same 0%, different 100%; To-nucleus distance distribution: same 45.5%, different 54.5%; Nearest-neighbor distance distribution: same 72.7%, different 27.3%. (F) Comparison of median distances before and after latA treatment using Wilcoxon rank sum tests (n = 55 pairs). Inter-organelle distance distribution: smaller: 54.5%; same: 9.1%, larger 36.4%; To-nucleus distance distribution: smaller 27.3%, same 36.4%, larger 36.4%; nearest-neighbor distance distribution: smaller 9.1%, same 81.8%, larger 9.1%. (G) Comparison of lysosomal spatial distributions in a control cell (left panel) versus a cell treated with trehalose (right panel; 50 mM, 12 hours). Red: lysosomes. Blue: nuclei; Scale bars: 15 μm. (H) The three distance distributions in control cells (n = 7) versus trehalose treated cells (12 hours, 7 cells). (I) Comparison of distributions in control cells and cells treated with trehalose using two sample Kolmogorov-Smirnov tests. Normalized inter-organelle distance distribution (n = 49 pairs): different 100%, same 0%; normalized to-nucleus distance distribution: different 85.7%, same 24.3%; nearest-neighbor distance distribution: different 97.9%, same 2.1%. (J) Comparison of median distances in control cells and cells treated with trehalose using Wilcoxon rank sum tests (n = 49 pairs). Normalized inter-organelle distance distribution: smaller 91.8%; same 0%; larger 8.2%; To-nucleus distance distribution: smaller 69.4%, same 16.3%, larger 14.3%; nearest-neighbor distance distribution: smaller 91.8%, same 8.2%, larger 0%.

Next, we investigated the role of the actin cytoskeleton in maintaining the spatial distribution of lysosomes. To this end, we compared the three distance distributions of lysosomes right before and 15 minutes after the treatment (Fig. 4D). We found that depolymerization of the actin cytoskeleton caused significant changes to the normalized inter-organelle distribution, the normalized to-nucleus distribution, and the nearest-neighbor distribution in 100%, ∽55%, and ∽25% of the cells, respectively (Fig. 4E; Table S4). In addition, we found that depolymerization of the actin cytoskeleton did not result in consistent increase or decrease in the median distances of lysosomes relative to each other and to the cell nucleus (Fig. 4F; Table S4). We reason that this is because interaction with the actin cytoskeleton does not alter the balance between centrifugal and centripetal transport of lysosomes. Taken together, our results indicate that interaction with the actin cytoskeleton is required for maintaining the positioning of lysosomes relative to each other but plays a minor role in maintaining their positioning relative to the nucleus and their crowding.

### A stable level of lysosomal biogenesis and autophagy is required for maintaining a stable spatial distribution of lysosomes

We have thus far investigated the role of the cytoskeleton in mediating the population composition and spatial distribution of lysosomes. Upregulation or downregulation of lysosomal biogenesis and autophagy can also cause substantial changes to the spatial distribution of lysosomes (Pu et al., 2016, Jongsma et al., 2016, Korolchuk et al., 2011, Sardiello et al., 2009), but such changes have not been quantitatively characterized at the whole-cell scale. To investigate how altered level of lysosomal biogenesis and autophagy may influence the stable spatial distribution of lysosomes, we treated COS-7 cells with 50 mM of trehalose (Sarkar et al., 2007), which activates TFEB, the master regulator of lysosomal biogenesis and autophagy (Sardiello et al., 2009, Settembre et al., 2011). We found that treatment of trehalose substantially increased the fraction of lysosomes clustering in the perinuclear region (Fig. 4G) and significantly decreased the distances between lysosomes and between lysosomes and the cell nucleus (Fig. 4H-J; Table S5), in agreement with observations of previous studies (Sardiello et al., 2009, Settembre et al., 2011). The treatment also reduced the median nearest-neighbor distance by ∽29%, substantially stronger in effect than the perturbations to the cytoskeleton analyzed previously (Fig. 3F, Fig. 3I, Fig 4D). Together, these results show that maintaining a stable spatial distribution of lysosomes requires a stable level of lysosomal biogenesis and autophagy.

### Lysosomes form dynamic clusters throughout the intracellular space

We have shown that lysosomes are spatially organized at the whole-cell scale because their spatial distributions are nonrandom, stable, yet distinct in single cells. Here we further investigate how they are spatially organized. A common pattern in the spatial distributions of lysosomes is their formation of a cluster in the perinuclear region, i.e. the perinuclear cloud (Pu et al., 2016, Jongsma et al., 2016, Korolchuk et al., 2011), which is defined by its substantially higher spatial density of lysosomes than the density of its neighboring area. Formation of this perinuclear cluster raises the question of whether lysosomes form clusters in other intracellular regions. To answer this question, we calculated and plotted their spatial density over the entire intracellular space (Fig. 5A-B; Materials & Methods). The plots revealed that lysosomes formed clusters throughout the intracellular space, which could be identified visually by their elevated spatial densities (Fig. 5A-B; Supplementary Movie S5). To identify these clusters computationally in an automated fashion, we used the DBSCAN algorithm (Ester et al., 1996)(Materials & Methods). Overall, we found that the identified clusters were dynamic and underwent turnover activities such as merging, splitting, appearance, and disappearance (Fig. 5C and Fig S4A).

**Figure 5.**
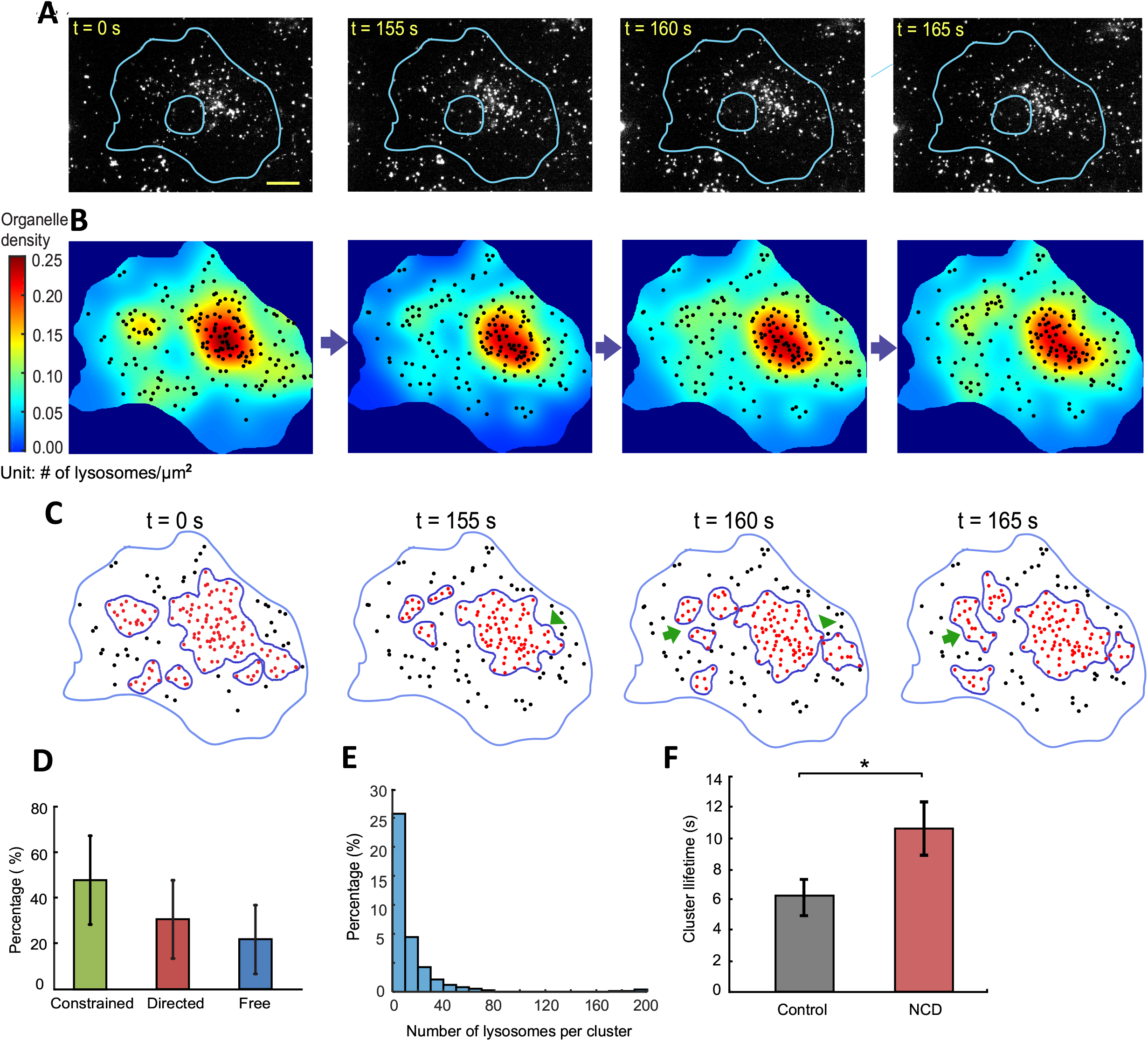
Lysosomes form dynamic clusters throughout the intracellular space. (A) Selected frames from a time-lapse movie of lysosomes labeled with dextran Alexa 488 in a COS-7 cell. Scale bar: 20 μm. (B) Color-coded spatial density plots of lysosomes calculated for the frames shown in (A). (C) Clusters of lysosomes identified computationally by DBSCAN. Arrow heads point to cluster splitting sites. Arrows point to cluster merging sites. (D) Composition of clusters, calculated for each cluster and then pooled for analysis. Constrained diffusion 47.9 ± 19.7%; directed transport, 30.6 ± 17.7%; free diffusion, 21.5 ± 14.9% (mean ± STD; n = 376 clusters from 9 cells, within 5 frames randomly selected from each cell). (E) Size distribution of clusters, measured in their numbers of lysosomes. Same clusters from same cells as those in (D). The average number of lysosomes was 15.4 ± 1.3 per cluster (mean ± SEM; n = 376 clusters from 9 cells). (F) Lifetime of large clusters with more than 10 lysosomes in the same cells before and after NCD treatment as shown in Figure 3D. Before treatment, 6.14 ± 1.22 seconds (mean ± SEM; n = 34 clusters from 8 cells); After NCD treatment, 10.69 ± 1.76 seconds (mean ± SEM; n = 24 clusters from 8 cells); p-value of one-tailed rank sum test, 0.013.

To quantitatively characterize the identified clusters, we measured some of their basic properties. First, we examined the composition of the clusters and found that, on avarage, 47.9% of their members were from lysosomes undergoing constrained diffusion, 30.6% from lysosomes undergoing directed movement, and 21.5% from lysosomes undergoing free diffusion (Fig. 5D). This composition was generally consistant with the composition of the entire lysosome population (Figure 3C) but was more heterogeneous given its larger variations (Figure 5D). Second, we quantified sizes of the clusters in terms of their numbers of lysosomes and areas. We found that the average number of lysosomes of the clusters was 15.4, with the largest cluster containing ∽200 lysosomes (Fig. 5E). The average area of the clusters was 47.2 µm^2^, with the largest cluster reaching ∽1084 µm^2^ (Fig. S4B). Lastly, we examined the lifetime of the clusters by following randomly selected clusters over time (Materials & Methods). We found that the average lifetime of the clusters was 10.9 ± 1.3 seconds (mean ±SEM; n = 30 clusters from 9 cells), with the longest lifetime reaching ∽20 seconds.

To determine the mechanisms underlying the formation and dynamic turnover of lysosomal clusters, we combined single particle tracking analysis (Fig. 3B-C) with clustering analysis of the lysosomes (Fig. S4C; Supplementary Movies S6 & S7). We found that the *de novo* formation of a new cluster at a certain location was mediated jointly by incoming lysosomes that undergo either directed movement or free diffusion together with lysosomes that undergo constrained diffusions at the location (Fig. S4C; Supplementary Movie S6). This finding is consistent with the measured properties of the clusters. The mean lifetime of a cluster is ∽11seconds. Within this time interval, the mean displacement of a lysosome undergoing constrained diffusion is less than 0.5 µm. Because the mean diameter of a cluster is 15 µm (Fig. S4B), formation of a new cluster requires long-range inward transport of lysosomes undergoing either directed movement or free diffusion. Because only a small fraction of lysosomes undergo free diffusion (Fig. 3C), we conclude that cluster formation is mediated primarily by lysosomes undergoing directed movement together with lysosomes undergoing constrained diffusion. Our single particle tracking provided no evidence that newly synthesized lysosomes appear in the clustering region. We should note that new clusters can also generate from event such as that splitting of an existing cluster and merging of existing clusters.

Consistent with our finding on cluster formation, we found that dynamic turnovers of lysosomal clusters were mediated primarily by long-range movements of lysosomes, especially those undergoing directed movement (Supplementary Movie S6). To further test this finding, we compared the lifetime of clusters in control cells versus cells treated with nocodazole (Fig. 3D). To minimize the influence of image noise on our life-time measurement, we focused on large clusters composed of more than 10 lysosomes. We found a significant increase in their mean lifetime by ∽73%, from 6.15 seconds in control cells to 10.68 seconds in nocodazole treated cells (Figure 5F), in support of our finding.

### Clustering of lysosomes substantially increases their interaction with late endosomes

That lysosomes form dynamic clusters throughout the intracellular space raises the important question of what functions these clusters may serve. Previous studies have assumed that the clustering of lysosomes in the perinuclear region when cells are under stress promote their interactions with partner organelles such as autophagosomes (Pu et al., 2016, Jongsma et al., 2016, Korolchuk et al., 2011). But this assumption has not been directly tested. Here we make a similar hypothesis, namely under normal conditions, clusters of lysosomes throughout the intracellular space increase interactions of lysosomes with partner organelles because of the increased lysosomal spatial density. To directly test this hypothesis, we imaged lysosomes and late endosomes concurrently at approximately 5 seconds per frame for 5 minutes (Fig. 6A; Supplementary Movie S8) and then analyzed their interactions. First, we confirmed that our experimental assay could reliably differentiate between lysosomes and late endosomes (Fig. S5; Materials & Methods). Then, we developed a computational method that identifies pairs of lysosomes and endosomes with a high likelihood of interacting with each other. Specifically, we identified a lysosome and an endosome as an interacting pair if they maintained a center distance below a threshold ranging from 400∽800nm for at least 25 seconds (Fig. 6B; Materials & Methods). Under the selected distance threshold and time threshold, we estimated that more than 80% of the detected interacting lysosome-endosome pairs had a spatial overlap of their fluorescence signals during most (> 80%) of the time they stayed within the threshold distance (Fig. S6A-E). Using our computational method, we counted interacting lysosome-endosome pairs in each cell and identified between 26 to 117 pairs on average per cell in 5 minutes (Fig. 6C-E). To examine the persistence of these candidate pairs, we calculated the time in which the pairs stayed within the distance threshold. We found that around ∽33% of the pairs stayed for more than 1 minute (Fig. 6F).

**Figure 6.**
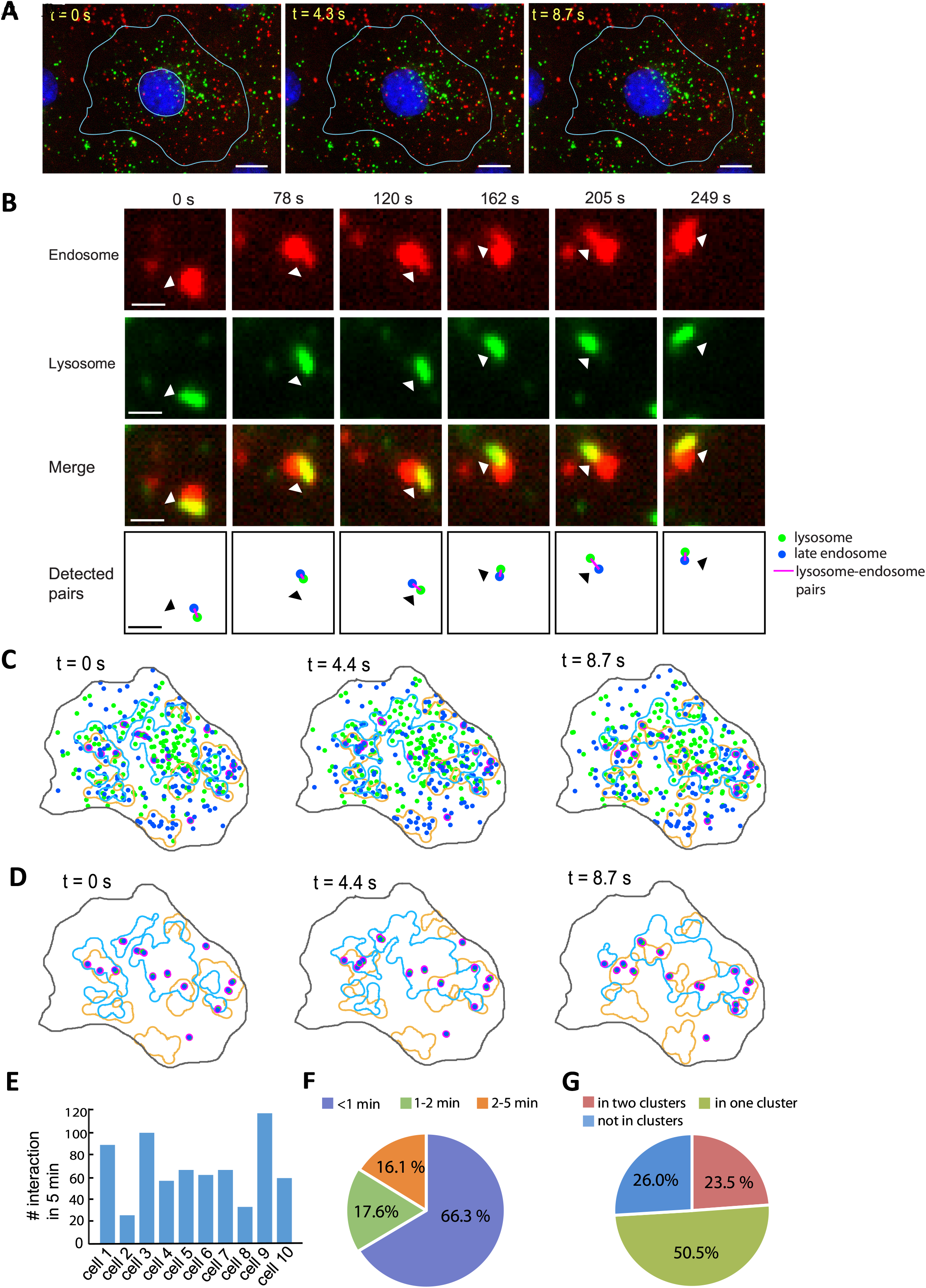
Clustering of lysosomes increases their interaction with late endosomes. (A) Selected frames from a time-lapse movie of a COS-7 cell in which lysosomes and late endosomes were labeled with dextran 488 and dextran Alexa 594, respectively. Scale bars: 15μm. (B) An example of computationally detected interacting lysosome-endosome pair that remained together for more than 4 minutes. Top three rows showed the actual fluorescence signals. Bottom row: computational detection result. Scale bars: 1 μm. (C) Simultaneous illustration of clusters of late endosomes and lysosomes detected from the movie in (A). Magenta: interacting lysosome-endosome pairs. Green: lysosome. Blue: late endosome. Light blue lines: lysosome clusters. Orange lines: endosome clusters. (D) Same as (E) but showing only the clusters and detected lysosome-endosome pairs. (E) Number of detected interacting lysosome-endosome pairs in 10 cells within 5 minutes: 89, 26, 101, 54, 64, 61, 63, 32, 117, 58. (F) Duration distribution of the detected interacting pairs staying together. (G) Relation between the location of interacting pairs and clusters of endosomes and lysosomes.

We then investigated the relations between the clusters of lysosomes and endosomes and the detected lysosome-endosome pairs. First, we identified clusters of lysosomes and clusters of endosomes, respectively (Fig. 6C). Then, we checked the distribution of the interacting lysosome-endosome pairs within and outside of the clusters. We found that 23.5% of the interacting pairs resided within areas in which lysosome clusters overlap with endosome clusters (Fig. 6G), 50.5% of the interacting pairs resided in either a lysosomal cluster or an endosomal cluster. In total, 74% of the interacting pairs were associated with at least one cluster. In comparison, 26.0% of the interacting pairs were not within any clusters. Together, these results suggest that clustering of lysosomes and endosomes substantially increased their interactions with endosomes.

To better understand the behavior of the computationally detected interacting lysosome-endosome pairs, we followed their activities visually. As an example, we found that a pair stayed together for 566 seconds to complete their fusion (Fig. 7A-B), consistent with the duration of fusion events reported in previous studies (Bright et al., 2005). Their fusion was confirmed based on their content exchange, as indicated by the fluorescence signals (Fig. 7A-B). This provides us an assay to further characterize our computational detection of interacting lysosome-endosome pairs. Given that the actual time required to complete the fusion is much longer than the mean lifetime of a lysosome cluster, at ∽11 seconds, we propose that clustering of lysosomes and endosomes promotes initiation of their interactions.

**Figure 7.**
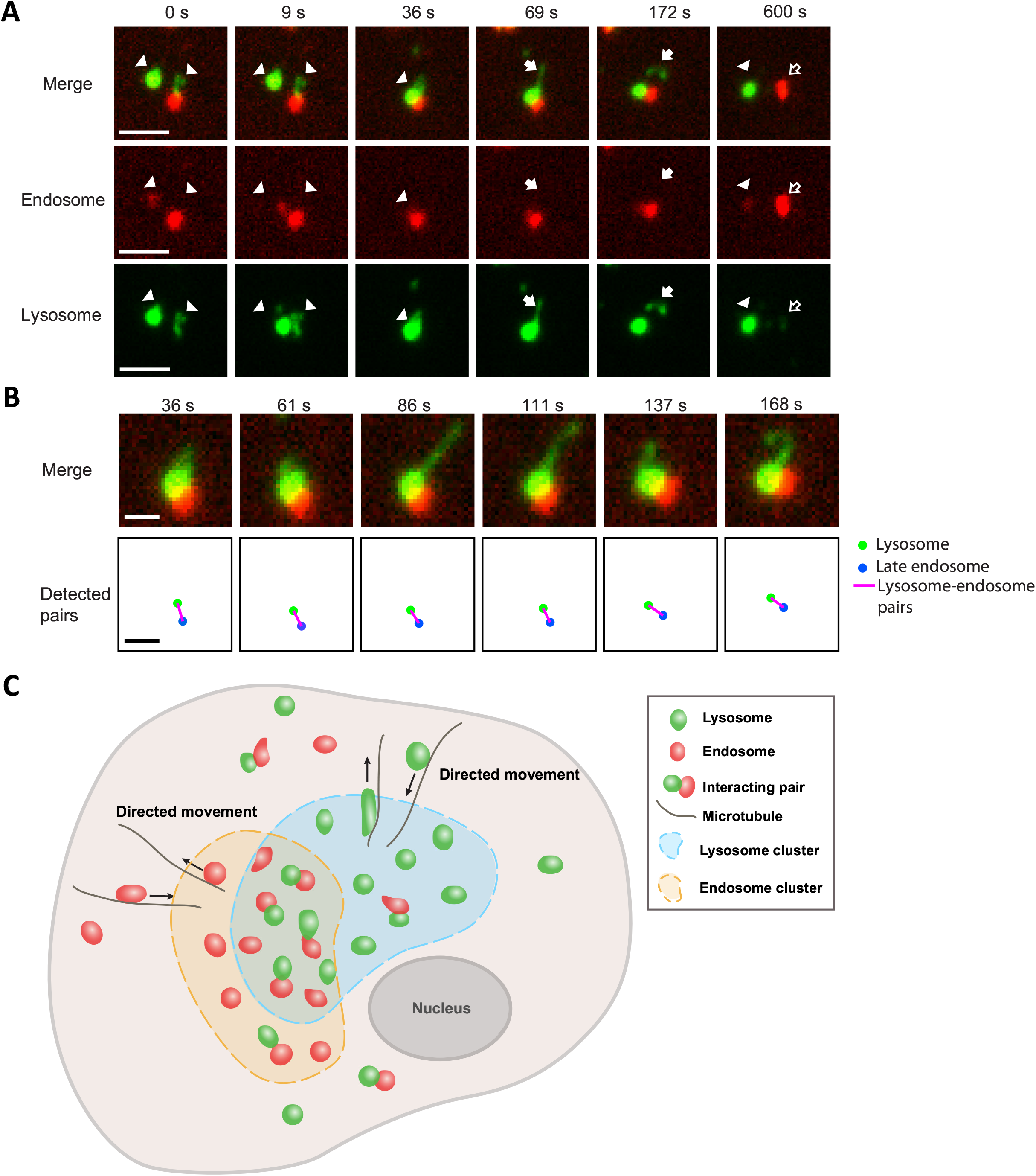
Tracking of fusion between lysosomes and endosomes and a model of lysosomal clustering. (A) An example of lysosome-endosome fusion in a COS-7 cell. Lysosomes (green) were labelled with dextran Alexa 488. Endosomes (red) were labelled with dextran Alexa 594. At 0s – 9s, two lysosomes (arrow heads) moved close to an endosome. At 36 s, the three organelles overlapped with each other in their fluorescence signals, presumably undergoing fusion or partial content exchange. This was followed by formation of a new lysosome at 69s-157 s indicated by arrows. At 600 s, separation and content exchange were evident given that the lysosome content (green fluorescence) was present in the endosome (hollow arrows) and vice versa. Arrow head: a lysosome. Solid arrow: a newly formed lysosome. Hollow arrow: an endosome that gained lysosomal content. Scale bar, 2.5μm. (B) Computational detection of lysosome-endosome interaction in the example shown in (A). Upper row: the actual lysosome-endosome pair. Bottom row: computational detection result. Frame rate during imaging: 4.2 second per frame. Scale bars, 1 μm. (C) A cartoon illustration of our active transport mediated clustering model.

## DISCUSSION

Clustering of lysosomes in the perinuclear region has been noted from early on in studies of lysosomal motility and positioning (Matteoni and Kreis, 1987). Currently, however, the predominant view is that individual lysosomes act independently and interact with partner organelles in an entirely random fashion. In this study we challenge this view. By combining high-resolution image analysis with spatial statistical analysis, we have found patterns in collective behavior of lysosomes. Specifically, we find that, despite their extensive long-distance movement, lysosomes maintain stable yet distinct spatial distributions at the whole-cell scale (Fig. 1F-H; Fig. 2A-C). Furthermore, we find that by forming dynamic clusters throughout the intracellular space, individual lysosomes work together to increase their spatial density locally and to promote their interactions with partner organelles such as endosomes (Fig. 5A-C; Fig. 6C-D). Lysosomes likely bear similarities with other intracellular organelles such as endosomes and peroxisome in their spatial organization because of their common evolutionary origins (Diekmann and Pereira-Leal, 2013) and their extensive interactions (Valm et al., 2017, Bonifacino and Neefjes, 2017, Luzio et al., 2007). Our findings that lysosomes maintain stable spatial distributions and form dynamic clusters develop new and potentially fundamental concepts in lysosomal biology, and organelle biology in general. They provide new insights into how organelle interactions are mediated and regulated at the whole-cell scale.

### Functional implications of stable yet distinct spatial distributions of lysosomes

Because of the crucial role of lysosomes in many important cellular processes, homeostasis of their functions is essential to cell homeostasis. We now know that at least some functions of lysosomes depend on their positioning (Johnson et al., 2016, Korolchuk et al., 2011). Because lysosomes maintain a stable spatial distribution at the whole-cell scale, homeostasis of their functions does not depend on fixed positions of individual lysosomes. While the spatial distribution of lysosomes in a single cell is stable, different cells maintain distinct distributions, even those with similar cell shapes (Fig. 2A-C). Single cells exhibit heterogeneity in many of their important properties (Altschuler and Wu, 2010). Here, our study reveals heterogeneity of single cells in their lysosomal spatial distributions.

### Mechanisms underlying stable spatial distributions of lysosomes

Our single particle tracking reveals that lysosomes in a single cell form different subpopulations (Fig. 3B-C). The subpopulation undergoing directed movement is maintained primarily by microtubule-based active transport. However, the subpopulation undergoing constrained diffusion is maintained only in part by interaction of lysosomes with the actin cytoskeleton (Fig. 4C). Interaction of lysosomes with ER(Bonifacino and Neefjes, 2017, Jongsma et al., 2016) and cytoplasmic crowding (Weiss et al., 2004) are possible additional mechanisms to be further investigated.

Overall, our analysis identifies three mechanisms whose coordinated effects mediate the stable spatial distribution of lysosomes at the whole-cell scale. The first mechanism is the balance between the subpopulation undergoing directed movement and the subpopulation undergoing constrained diffusion. This balance is essential to maintaining a stable composition of the lysosomal population. The second mechanism is the balance between microtubule-based transport in the centrifugal direction and the centripetal direction. This balance is essential to maintaining the overall positioning of lysosomes relative to each other and to the nucleus. The third mechanism is homeostasis in the biogenesis of lysosomes, which is essential to maintaining the level of crowding between lysosomes (Fig. 4H-J). Additional mechanisms, especially upstream regulatory mechanisms, almost certainly are involved in maintaining the stable spatial distribution of lysosomes.

### Functional implications of clustering of lysosomes

Using spatial statistical analysis techniques, we find that clustering of lysosomes is not restricted to the perinuclear region. Instead, lysosomes form dynamic clusters throughout the intracellular space (Fig. 5C). Furthermore, using single particle tracking, we directly show that clustering of lysosomes promotes their interactions with late endosomes (Fig. 6C-D).

A key benefit of the dynamic clustering of lysosomes is that it allows tuning of interaction of lysosomes with partner organelles locally throughout the intracellular space. By increasing spatial density of lysosomes locally, formation of clusters increases the likelihood of their collisions and interaction with partner organelles such as endosomes. Importantly, dynamic turnover of the clusters makes it possible to switch on or off such effect. Given the common evolutionary origins (Diekmann and Pereira-Leal, 2013) and extensive interactions (Valm et al., 2017, Bonifacino and Neefjes, 2017, Luzio et al., 2007) of lysosomes with other organelles, we speculate that dynamic clustering is general mechanism for promoting and regulating organelle interactions.

Another key benefit of the dynamic clustering of lysosomes is that it facilitates the maintenance of lysosomal homeostasis. Lysosomes contain high levels of digestive enzymes (Xu and Ren, 2015). Rupture of lysosomal membrane under oxidative stress releases the enzymes into the cytoplasm and is known to trigger apoptosis under many conditions (Colletti et al., 2012, Sun et al., 2010, Kanzawa et al., 2004, Werneburg et al., 2002). Therefore, to promote interactions of lysosomes with partner organelles through global crowding via nonspecific increase in their total number is not preferable, not only because this strategy lacks spatial specificity but also because it is detrimental to cell physiology. Dynamic clustering of lysosomes promotes their interactions with partner organelles without the need to increase their total numbers and therefore facilitates the maintenance of lysosomal homeostasis.

### A model of lysosomal clustering

To summarize our findings on lysosomal clustering, we propose what we refer to as an active transport mediated clustering model (Fig. 7C). In particular, we propose that formation and dispersion of lysosomal clusters can be controlled by controlling the active transport of lysosomes.

## MATERIALS AND METHODS

### Cell culture and organelle labeling

COS-7 were maintained in Dulbecco’s modified Eagle medium (DMEM) supplemented with 10% fetal bovine serum. BS-C-1 cells were maintained in Eagle’s Minimum Essential Medium (EMEM) supplemented with 10% fetal bovine serum. Cells, culture media, and serum were purchased from American Type Culture Collection (Manassas, VA).

Lysosomes in COS-7 cells were labeled by transfecting 200-300 ng LAMP1-mCherry. Transfection of COS-7 cells was performed using the Neon electroporation system (Invitrogen, Carlsbad, CA). Briefly, 2×10^5^ cells were suspended in a 10 µl pipette tip and electroprated under a pulse voltage of 950 V, a pulse width of 30 ms, and a pulse number of 2. Following transfection, cells were seeded at 6.7×10^4^ per 20 mm glass well in 35 mm dishes (MatTek, Ashland, MA) and incubated for 24-48 hours before imaging. Lysosomes in BS-C-1 cells were labeled as described in (Humphries et al., 2011, Kilpatrick et al., 2015, Bright et al., 2005). Briefly, cells were incubated with 0.5 mg/ml Dextran Alexa 488, 10000 MW (Molecular Probes, Eugene, OR) for 3-4 hours followed by 16 hours of chasing. Late endosomes and lysosomes in COS-7 cells were co-labelled as described in (Bright et al., 2015, Bright et al., 2005). Briefly, cells were incubated with 0.5 mg/ml Dextran Alexa 488 for 3 hours, followed by 20-27 hours of chasing to mark lysosomes. Then cells were incubated with 0.5 mg/ml Dextran Alexa 594, 10000 MW (Molecular Probes, Eugene, OR) for 10 minutes followed by 10 minutes of chasing to mark endosomes before imaging. That the co-labeling scheme reliably differentiated late endosomes from lysosomes was validated as described in (Luzio et al., 2007, Bright et al., 2005) by immunostaining of Mannose-6 phosphate receptor (M6PR), a marker present in endosomes but not in lysosomes, using a monoclonal M6PR antibody (MA1-066; Thermo Fisher).

### Drug treatment

To depolymerize microtubules, COS-7 cells were treated with 2.5 µM nocodazole (Sigma-Aldrich, St. Louis, MO) and imaged before and 15 minutes and 30 minutes after treatment for the same cells. To depolymerize actin filaments, COS-7 cells were treated with 800 nM latrunculin A (Sigma-Aldrich, St. Louis, MO) and imaged before and at 7-8 minutes and 15-16 minutes after treatment. To examine the effect of latrunculin A treatment on the actin cytoskeleton, cells were fixed and stained with fluorescent phalloidin (Actin-staining 488 fluorescent phalloidin; Cytoskeleton, Denver, CO) following instructions of the manufacturer. To upregulate lysosomal biogenesis and autophagy, cells were treated with 50 mM trehalose (Sigma-Aldrich, St. Louis, MO) as described in (Sarkar et al., 2007, Porter et al., 2013, Palmieri et al., 2017) and imaged after 12 hours of treatment. To inhibit cytoplasmic dynein, COS-7 cells were treated with 80 µM ciliobrevin D (Sigma-Aldrich, St. Louis, MO) and imaged before and 1 hour after treatment for the same cells.

### Controlling cell shape using patterned protein substrates

Cells were grown into defined shapes by culturing them on patterned protein substrates made by micro-contact printing (Azioune et al., 2009, Xia and Whitesides, 1998, Singhvi et al., 1994)

*PDMS stamp fabrication*: Protein substrate patterns were designed using AutoCAD (Autodesk, San Rafael, CA). A plastic mask with designed circular substrate patterns (68 µm in diameter) was produced by CAD/ART Services (Bandon, OR) at 10-µm resolution. Master molds were fabricated by spin coating a 4-µm thick layer of SPR 220-3.0 (MicroChem, MA) onto a 2 µm thick coverslip glass (Fisher Scientific, Hampton, NH) followed by UV exposure at 365 nm using a custom made UV illumination system. Polydimethylsiloxane (PDMS) stamps were made by mixing Sylgard 184 (Dow Corning, MI, US) PDMS base with curing agent at the ratio of 1:10, followed by 1 hour defoaming under vacuum and curing for 12-24 hours at 65 °C. Stamps approximately 1 cm × 1 cm in size were cut from the PDMS blocks for micro contact printing (Azioune et al., 2009).

*Micro contact printing of protein substrates*: PDMS stamps were sonicated in 75% ethanol for 30 minutes and dried by nitrogen blowing under a laminar hood. The stamps were then coated with 200 µL of 20 µg/ml fibronectin (Thermo Fisher Scientific, Waltham, MA) in PBS and incubated at room temperature for 1 hour. Alex 594 conjugated fibrinogen (Molecular Probes, Eugene, OR) was added to the fibronectin solution at a final concentration of 8 µg/ml for visualization of printed patterns (Azioune et al., 2009). The stamps were rinsed in deionized water and dried under a laminar hood. The stamps were then placed with the pattern side down on glass surfaces in 35 mm MatTek dishes (Ashland, MA) with 2.0 cm wells. After 1 hour the stamps were removed to release patterned proteins (Palchesko et al., 2012). To prevent cell attachment in unpatterned area, the printed glass surfaces were coated with 0.1 mg/ml of PLL-g-PEG (Surface Solutions, Dübendorf, Switzerland) in PBS for 1 hour and rinsed with PBS for three times.

*Cell culture on patterned protein substrates*: Dextran-488 labeled BS-C-1 cells were trypsinized 16 hours after labeling and seeded onto patterned substrates. Unattached cells were removed 45-60 min after seeding. Imaging was performed 4-7 hours after seeding.

### Immunofluorescence

Cells were grown in 35mm dishes (MatTek, Ashland, MA) and fixed with 4 % formaldehyde for 8 minutes and permeablized in 0.2% Triton X-100/PBS for 5 minutes at room temperature. Cells were then blocked with 5% normal goat serum for 1 hour and incubated with primary antibody at 4°C overnight and secondary antibodies at room temperature for 1 hour. After each antibody incubation step, cells were washed five times with DPBS with Ca^2+^ and Mg^2+^, 5 minute each time. Nuclei were then labeled with Hoechst 33342 (Sigma-Aldrich). Samples were imaged in DPBS with Ca^2+^ and Mg^2+^. To differentiate lysosomes from late endosomes, mouse monoclonal antibody against mannose 6-phosphate receptor (1:500, Thermo Fisher MA1-066) was used. The secondary antibody used was Alexa Fluor 594-conjugated goat anti-rabbit IgG (1:400, Abcam ab150116, Cambridge, UK).

### Fluorescence microscopy

Imaging was performed on a Nikon Eclipse Ti-E inverted microscope with a CoolSNAP HQ2 camera (Photometric) and a 100×/1.40 NA or 60×/1.40 NA oil objective lens. The effective pixel sizes were 0.0645μm at 100× and 0.107 μm at 60×, respectively. For ciliobrevin treatment, imaging was performed with a Zyla (Andor) CMOS camera with a 60×/1.40 NA oil objective lens. The effective pixel sizes was 0.11 μm at 60×. During live imaging, cells were maintained in a Tokai HIT stage incubator (Tokai, Shizuoka, Japan) at 37 °C with 5% CO2. LAMP1-mCherry for labeling late endosomes and lysosomes and Dextran Alexa-594 for labeling endosomes were imaged using a TRITC filter set. Dextran Alexa-488 for labeling lysosomes was imaged using a FITC filter set. For each data set, at least six cells from two to three independent experiments were imaged.

Lysosomes in non-patterned cells were imaged at 4 frames per second. Lysosomes in patterned cells were imaged at 2.5 frames per second. One hour movies of lysosomes in COS-7 cells were imaged at 25 seconds per frame. In latrunculin A, nocodazole, and ciliobrevin D treatment experiments, cells were imaged at 10 frames per second. In endosome-lysosome interaction experiments, cells were imaged at 4.2-5 seconds per frame.

### Detecting and tracking lysosomes in images

To identify lysosomes in a given image, the Spot Detector plugin of the Icy software (de Chaumont et al., 2012) was used. Cartesian coordinates of detected lysosomes were exported into an Excel XLS file and then imported into our custom MATLAB (MathWorks, Natick, MA) software for further spatial statistical analysis of their distributions. To track movement of lysosomes in a given time-lapse movie, the Spot Tracking plugin of Icy (de Chaumont et al., 2012, Chenouard et al., 2013) was used. Similar as in lysosome detection, recovered lysosome trajectories were exported into an Excel XLS file and then imported into our custom software for further mean-square-displacement (MSD) analysis and spatial statistical analysis. Our software is openly available in source code at https://github.com/ccdlcmu/LysosomeSpatialOrganization_code.

### Complete spatial randomness test of lysosomal distributions

As an essential step in statistical analysis of spatial point processes (Illian et al., 2008), complete spatial randomness (CSR) test was performed on the distribution of lysosomes at the whole-cell scale to check whether it was entirely random. Specifically, for a given cell, its boundary was manually traced using the *imfreehand* function in MATLAB. The boundary geometry data and the coordinates of detected lysosomes were then passed to *R* for calculation of the Ripley’s *K*-function by calling the *spatstat* package. To provide a reference for comparison, a homogeneous Poisson point process was simulated in the same cell geometry with the mean number of simulated lysosomes matching the number of actual lysosomes. The *K*-function was calculated from 99 rounds of simulation. Because the *K*-function for a homogeneous Poisson process has the form *K*(*r*) = *πr*^2^ (Illian et al., 2008), we subtracted *πr*^2^ from the calculated *K*-functions of both actual and simulated lysosomal distributions for convenience of comparison (Fig. 1C-E). Substantial separation of the actual *K*-function from the reference would indicate non-randomness in distribution.

### Classification of lysosomes based on their modes of movement

Trajectories of lysosomes were recovered through single particle tracking as described above. From each trajectory that lasts at least 25 frames, which correspond to 2.4 seconds in imaging, the mean square displacement *(MSD)* function was calculated as described in (Saxton, 1997) with the maximum lag of 24 frames. Because *MSD* is a function of time, we assume the following simplified model:

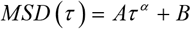

in which *α* can be used to classify different modes of movement (Qian et al., 1991). To determine *α*, the *MSD* was fitted using the MATLAB function *nlmfit*. The mode of movement of the corresponding lysosome was then classified according to the following table

| $\alpha$ | Mode of Movement |
| --- | --- |
| $\alpha < 0.9$ | Constrained diffusion |
| $0.9 < \alpha < 1.2$ | Free diffusion |
| $\alpha > 1.2$ | Directed movement |

## Characterization of lysosome distribution at the whole-cell scale

Lysosome distribution at the whole-cell scale was characterized using statistical distributions of distances between individual lysosomes as well as distances between individual lysosomes and the cell nucleus. These distributions were constructed based on the point process theory of spatial statistical analysis (Illian et al., 2008, Diggle, 2014) and were calculated from detected lysosome positions using our custom software.

*(Normalized) Inter-organelle distance distribution:* For a given cell, the distribution is calculated from all pairwise distances of its lysosomes (Fig. S1A), which characterizes spacing between lysosomes at the whole-cell scale. Specifically, for the *i^th^* and *j^th^* lysosomes, whose positions are denoted 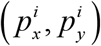 and 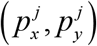, respectively, their inter-organelle distance *D_IO_*(*i,j*) is their Euclidean distance, 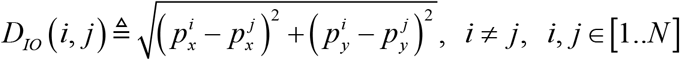, where *N* is the total number of lysosomes. To account for variations in cell sizes, the normalized inter-organelle distance 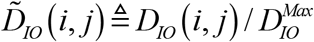 is used, where 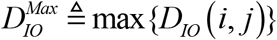 is the maximum inter-organelle distance. For each cell, after calculating its complete set of non-normalized inter-organelle distances, 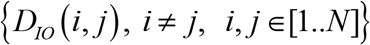, or normalized inter-organelle distances, 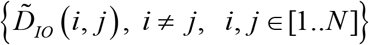, the probability density function (pdf) of the distances is estimated using MATLAB function *ksdensity* with default parameters, including a normal kernel whose bandwidth is optimized for density estimation.

*(Normalized)To-nucleus distance distribution:* For analysis given cell, the boundary of its DAPI-stained nucleus is manually traced and then approximated by the convex hull of the set of boundary points, We denote the set of boundary points as *S(C).* For the *i^th^* lysosome, its to- nucleus distance *D_TN_(i)* is its shortest distance it to the nucleus convex hull, defined as 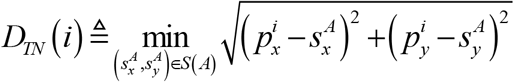. The normalized to-nucleus distance 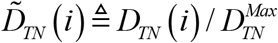 is used to account for variations in cell sizes, where 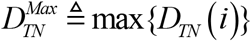 is the maximum to-nucleus distance. For each cell, the probability density function of its complete set of non-normalized to-nucleus distances, 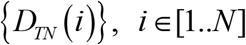 or normalized to-nucleus distances, 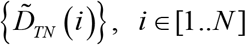 is estimated using MATLAB function *ksdensity* with the same parameter setting as for the inter-organelle distance distribution.

*Nearest-neighbor distance distribution:* For a given cell, this distribution is calculated from all nearest neighbor distances of its lysosomes. It characterizes the shortest distances between individual lysosomes at the whole-cell level. Specifically, for the *i^th^* lysosome, its nearest-neighbor distance *D_NN_(i)* is defined as 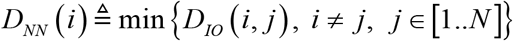. The probability density function of the complete set of nearest-neighbor distances, 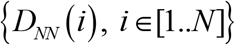 is estimated using MATLAB function *ksdensity* with the same parameter setting as for the inter-organelle distance distribution.

## Quantification of differences in lysosome distributions at the whole-cell level

After defining the distance distributions for characterizing lysosome spatial distributions at the whole-cell level, it is often necessary to compare such distributions at two different time points in the same cell or between two different cells. To compare two distance distributions, represented by *pdf_i_* and *pdf_j_*, respectively, we quantified their difference using the following intersection measure (Cha, 2007), also referred to as the Sorensen distance, which quantifies the level of non-overlap between the two distributions.

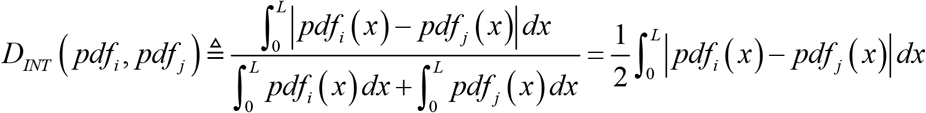

For normalized distance distributions, *L* equals 1. For non-normalized distance distributions, *L* equals the larger one of the two maximal distances of the two distributions.

## Estimating spatial density distributions of lysosomes

The spatial density distribution of lysosomes within a single cell represent the number of lysosomes per unit area at different locations inside that cell. This distribution was estimated using the R package *spatstat* (Baddeley and Turner, 2005, Baddeley et al., 2016). Specifically, for a given cell, with its size measured in micrometers and its shape represented by a polygon, a window of the same size and shape was created using the R function *owin*, Lysosomes detected within the cell as described above were used to create a point pattern object using the R function *ppp*. The spatial density distribution was then estimated using a kernel-based method by calling the R function *density.ppp.* The estimated spatial density distribution 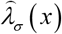 is defined by the following equation:

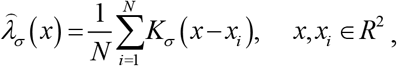

where *K*_σ_(·) is an isotropic 2D Gaussian kernel whose standard deviation σ is also referred to as the bandwidth of estimation, *x* represents any given position within the cell, *x_i_* is the position of the *i*^th^ lysosome, and *N* is the total number of lysosomes in the cell. The grid size for estimating the spatial density distribution was set to be 1 µm. The grid number was determined by the size of the smallest rectangle that circumscribes the cell. Note that the density distribution estimated by *spatstat* is not in the form of a probability density function. Instead, it directly represents the number of lysosomes per unit area and is thus convenient for interpretation.

The estimation bandwidth σ was chosen using the R function *bw.diggle* by minimizing the means quared error of the density estimation (Berman and Diggle, 1989). For estimating spatial density distributions of lysosomes over time in a time-lapse movie, σ was chosen based on the first frame and then kept the same for all subsequent frames.

## Density-based clustering of lysosomes

To study collective behavior of lysosomes, clusters of lysosomes were identified using a density based clustering algorithm DBSCAN (Ester et al., 1996). Briefly, a lysosome was randomly selected as a seed. The algorithm then searched a circular neighborhood of radius *r* centered at the seed. If the total number of lysosomes in the neighborhood was less than a preset threshold *k,* the seed was excluded as a noise point. If the number equaled or exceeded the threshold *k*, the seed lysosome with its neighbors were considered to be in a high density region and incorporated in a cluster. The incorporated neighboring lysosomes were then set as new seeds. This process was repeated until the number of lysosomes in a new neighborhood fell below threshold *k.* The algorithm repeated this process for the rest of lysosomes not in any cluster. Note that in this way, each cluster has at least *k* lysosomes within a neighborhood of radius *r.* A threshold setting of *k* ≥ 4 was recommended for good performance and reasonable computational cost (Ester et al., 1996). We chose a threshold of *k* = 5 to balance stringency of thresholding and sensitivity for identifying small clusters.

To set the neighborhood radius *r* for a given threshold *k,* the spatial densities of lysosomes should be higher than the spatial densities of a random and uniform distribution of lysosomes with the same total number of lysosomes. This distribution was determined through computer simulation. Specifically, for a given cell, 99 simulations were performed using the *spatstat* package function *runifpoint,* which used the rejection method (Illian et al., 2008) to generate a random and uniform point pattern inside the cell geometry, with the number of simulated lysosomes matching the actual number of lysosomes in the cell. The spatial density distribution of the simulated point pattern was then computed using the *spatstat* function density.ppp.

A threshold spatial density, denoted *λ_thresh_* was then set to be 95% of the maximal spatial density from simulation. The search radius was then calculated as 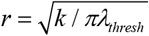, with a threshold of *k* = 5. For a time-lapse movie, the simulation was performed based on the number of lysosomes in the first frame. The search radius determined was then used for the rest of the frames.

## Analyzing interactions between lysosomes and partner organelles

*Nearest-neighbor distribution between lysosomes and partner organelles:* The nearest neighbor distance distribution between lysosomes was extended to characterize the relative positioning of lysosomes with partner organelles, such as late endosomes. For each lysosome, its distance to the nearest partner organelle was calculated. Then the probability density function was estimated from the nearest neighbor distance of all lysosomes using MATLAB function *ksdensity and* was referred to as the nearest-neighbor distance distribution between lysosomes and the partner organelles.

*Detecting interacting pairs of organelles:* Candidates of interacting pairs were detected first based on spatial proximity. A pair of organelle was considered a candidate if they distance was smaller than a threshold distance *D*_min_, which was determined based the following formula

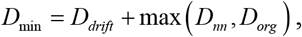

This threshold was calculated for each cell and typically ranged from 0.4 µm to 0.8 µm. *D_drift_* denotes the distances organelles travel within the lag of switching between channels during imaging. For the imaging setup of this study, the lag is roughly 1 s, and the corresponding drift was estimated to be ∽0.2 µm based on single particle tracking of late endosomes and lysosomes. The nearest neighbor distance *D_nn_* was chosen as the lower 10% quantile of the nearest neighbor distance between lysosomes and the partner organelles. Lastly, the diameters of organelles were also considered when *D_nn_* is smaller than the lower bound of organelle size *D_org_*, which can occur because the position of an organelle was represented by its centroid. The lower bound of organelle size *D_org_* was set to be 0.25 µm, given the diffraction limit of light microscopy and the fact that late endosomesand lysosomes are often larger larger than this size.

From the candidate pairs, interacting pairs were identified if the time they stay within the distance threshold is longer than a time threshold. This threshold is set to be 25 seconds based on a >85% quantile of the time different pairs of organelles staying with the distance threshold. The detection results are then visually inspected to exclude errors. Under the distance threshold and the time threshold selected, more than 80% of the detected interacting pairs of lysosomes and endosomes had a spatial overlap of their fluorescence signal during at least 80% of the time they stayed within the threshold distance to each other.

*Detecting interacting pairs within organelle clusters:* To detect interacting pairs inside organelle clusters, the MATLAB function *boundary* was used to determine boundaries of identified organelle clusters, represented as polygons. Then the function *inpolygon* was used to select interacting pairs. To detect interacting pairs inside overlaying regions of clusters of both types of organelles, endosomes in any interacting pair that lied within or on boundary of any lysosome cluster 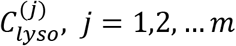, were detected as candidates. Then the candidate endosomes that fell within or on the boundaries of any endosome cluster(s) were selected. After this step, we accept the candidate pair if the interacting lysosomes also reside in the same endosome cluster boundary and the lysosome cluster boundary.

## Author Contributions

G Y., K.K., and Q.B. conceived and designed the project. Q.B. performed the experiments. G.Y., Q.B., and G.R. designed and performed the data analysis. K.K. contributed to experimental design and data analysis and provided reagents. G.Y. and Q.B. wrote the manuscript with contributions from all other authors.

## Acknowledgements

Q.B. acknowledges support of a Bertucci Graduate Research Fellowship. K.K. acknowledges support of NIH grants NS096755 and NS094860. G.Y. acknowledges support of NSF CAREER grant DBI-1149494.

## Supplemental Text and Figures

**Figure S1.**
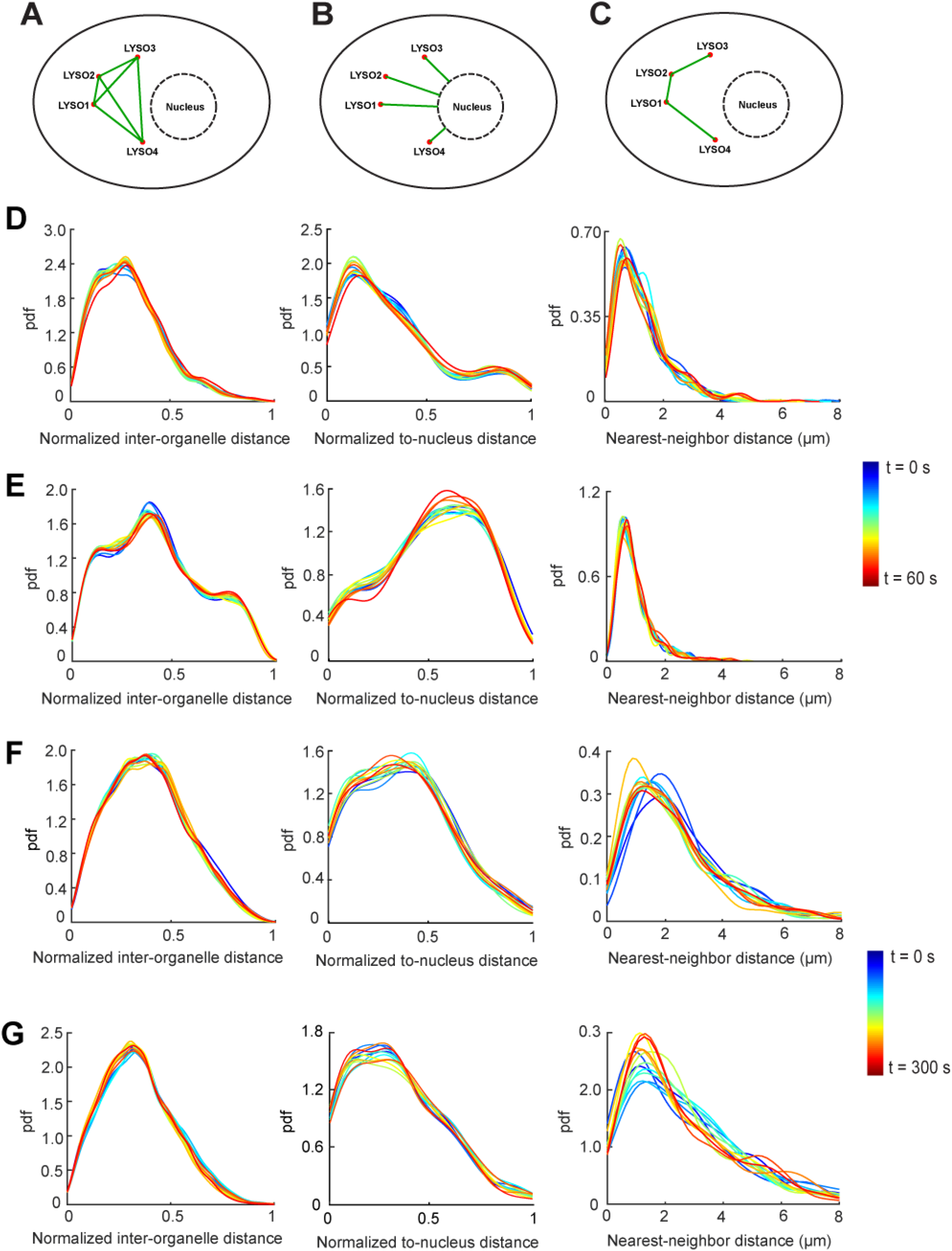
Spatial distributions of lysosomes remain stable over time in single cells. (A) A cartoon illustration of inter-organelle distances. (B) A cartoon illustration of to-nucleus distances. (C) A cartoon illustration of nearest-neighbor distances. (D) and (E): three distance distributions of lysosomes from two BS-C-1 cells. Each distribution was plotted every 5 seconds over 60 seconds, hence 13 plots. The temporal variations were quantified using Sorensen dissimilarity scores. A total of 13 distributions were compared pairwise, hence 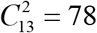 pairs. pdf: probability density function. (D) Temporal variations (mean ± STD; n = 78): normalized inter-organelle distance distribution, 1.77% ± 0.64%; normalized to-nucleus distance distribution, 2.53% ± 0.72%; nearest-neighbor distance distribution, 8.64% ± 2.65%. (E) Temporal variations (mean ± STD; n = 78): normalized inter-organelle distance distribution, 1.87% ± 0.61%; normalized to-nucleus distance distribution: 2.60% ± 1.04%; nearest-neighbor distance distribution, 6.07% ± 1.51%. (F) and (G): three distance distributions of lysosomes from two COS-7 cells. Each distribution was plotted every 25 seconds over 300 seconds, hence 13 plots. (F) Temporal variations (mean ± STD; n = 78): normalized inter-organelle distance distribution, 1.53% ± 1.43%; normalized to-nucleus distance distribution, 2.04% ± 1.75%; nearest-neighbor distance distribution, 5.19% ± 4.47%. (G) Temporal variations (mean ± STD; n = 78): normalized inter-organelle distance distribution, 1.71% ± 1.66%; normalized to-nucleus distance distribution, 1.94% ± 1.63%; nearest-neighbor distance distribution, 5.55% ± 4.49%.

**Figure S2.**
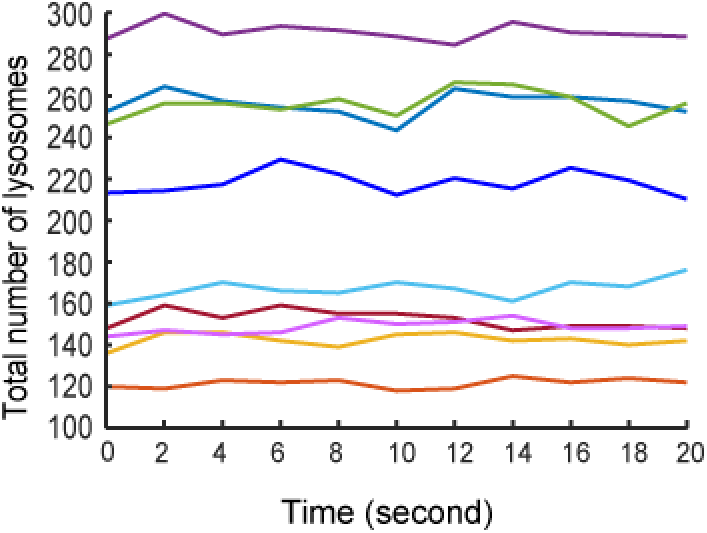
Total numbers of lysosomes remain stable in single cells. Total numbers of lysosomes in COS-7 cells during 20 seconds of imaging, plotted every 2 seconds for each cell, n = 9 cells. Frame rate: 10 frames per second. Total numbers of lysosomes in each cell (mean ± STD): 122 ± 2, 142 ± 3, 149 ± 3, 152 ± 4, 167 ± 5, 218 ± 6, 255 ± 7, 256 ± 6, 290 ± 4.

**Figure S3.**
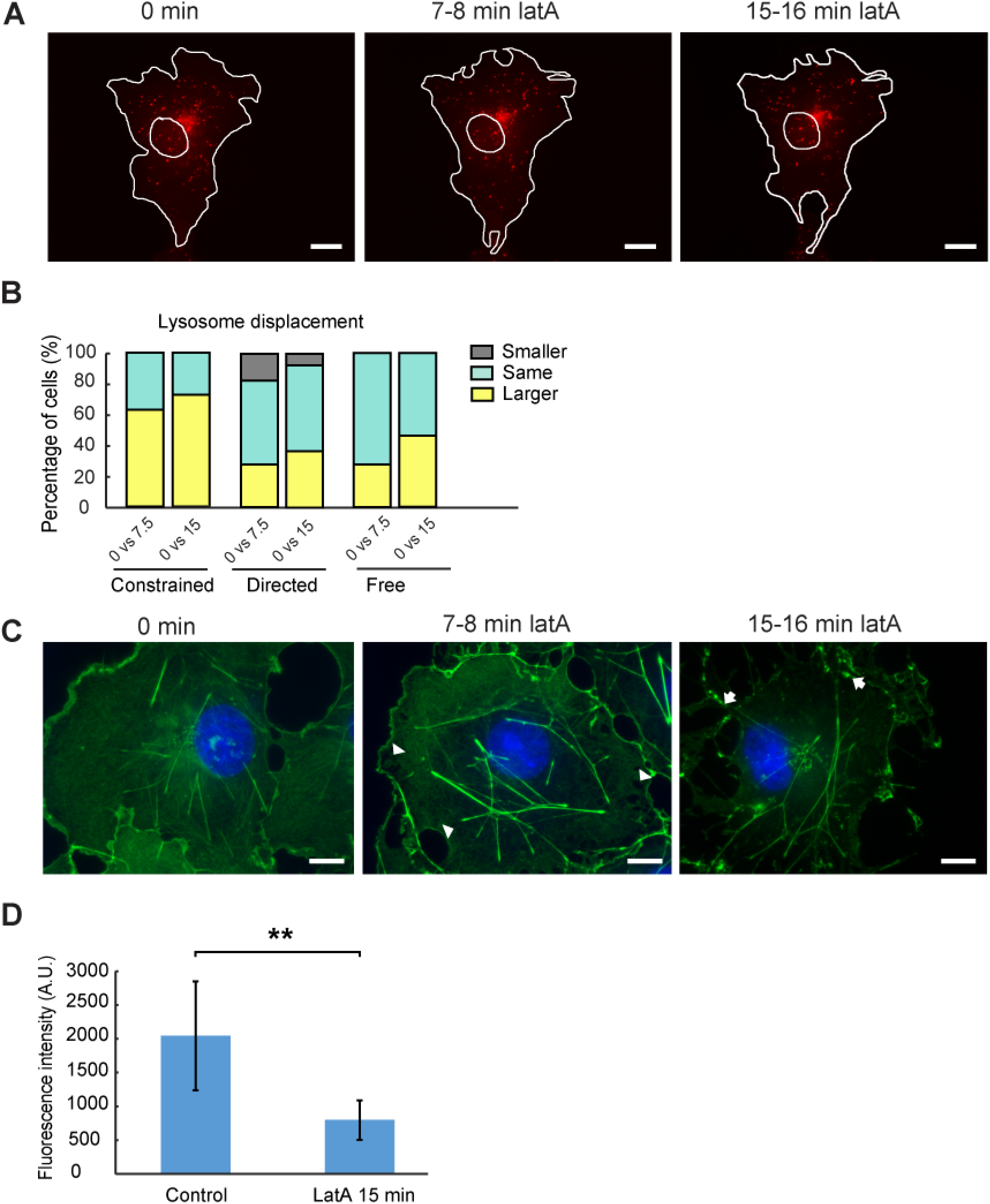
Changes of actin network and lysosomal movement upon latrunculin A treatment. (A) Shape changes of a COS-7 cell transfected with mCherry-Lamp1 at different time points before and after treatment of 0.8 μM of latrunculin A. Red, lysosomes. Scale bars: 15 μm. (B) Comparison of displacement of different lysosomal subpopulations within 5 seconds before and after latA treatment in single cells: before treatment (0 min) vs 7.5 min; before treatment (0 min) vs. 15 min. n = 11 cells. One-tailed t-tests were performed with a cutoff significance level of 0.05. Constrained, 0 min vs 7.5 min: smaller, 0%; same 36.4%; larger: 63.6%; constrained, 0 in vs 15 min: smaller 0%; same 27.3%; larger 72.7%. Directed, 0 vs 7.5 min: smaller 18.2%; same 54.5%; larger 27.3%; directed, 0 vs 15 min: smaller 9.1%, same 54.5%; larger 36.4%. Free diffusion, 0 vs 7.5 min: smaller 0%; same 72.7%; larger 27.3%; free diffusion, 0 vs 15 min: smaller 0%; same 54.5%, larger 45.5%. (C) COS-7 cells fixed and stained with phalloidin (Actin-stain 488) under control condition and after 7.5 min and 15 min of latA (0.8 μM) treatment. Arrow heads point to regions with reduced phalloidin fluorescence signals. Arrows point to actin patches. Scale bars, 10 μm. (D) Quantification of fluorescence intensity of actin in control and latA treated cells. A.U., arbitrary unit. Control: 2048 ± 807 (mean ± STD; n = 7 cells); latA 15 min: 799 ± 289 (mean ± STD; n = 7 cells). A one-tailed t-test was performed; p-value, 0.0098.

**Figure S4.**
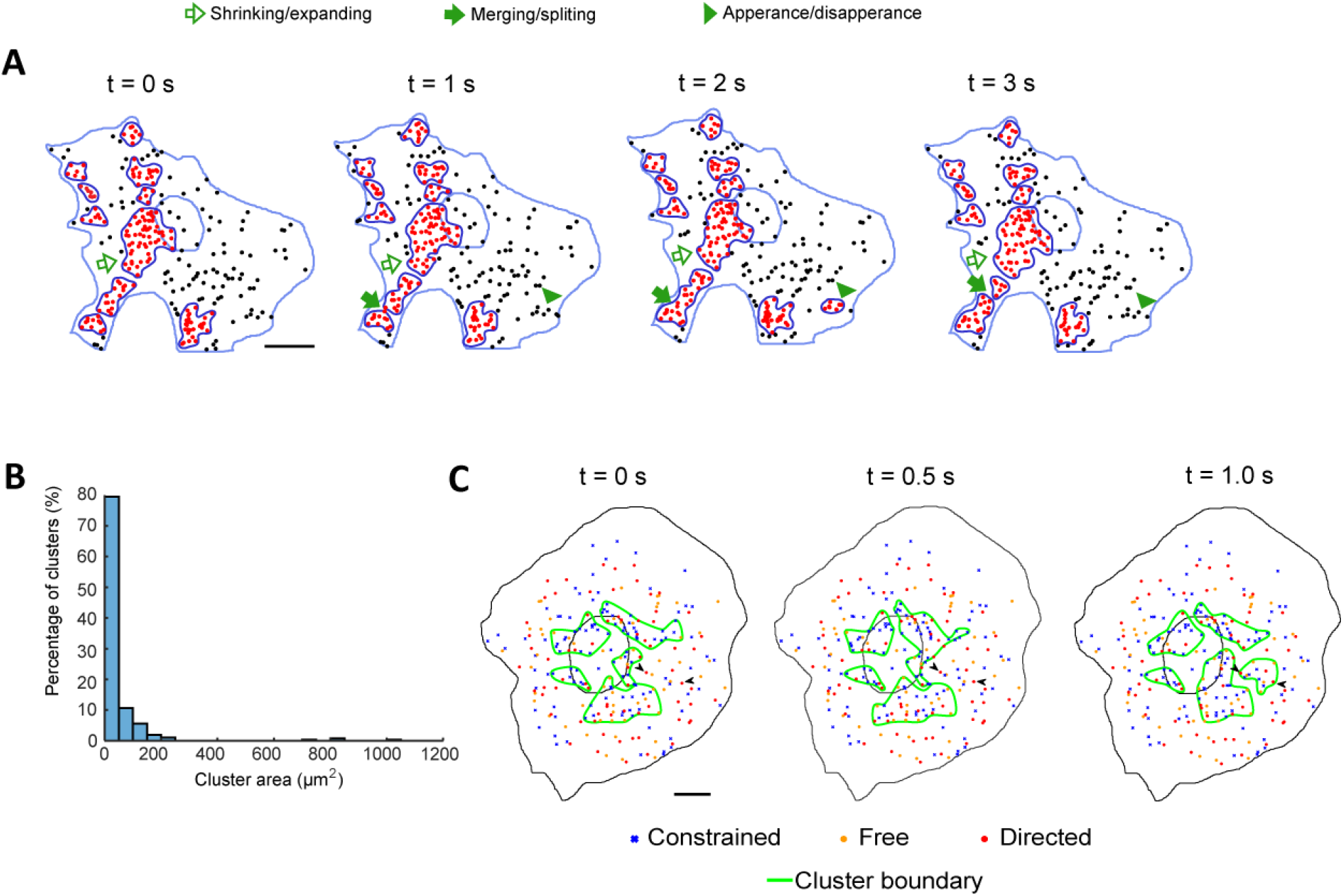
Dynamic turnover and size distribution of lysosomal clusters. (A) Various dynamic turnover events of lysosomal clusters in a COS-7 cell. Scale bar, 15 μm. (B) Size distribution of lysosomal clusters in COS-7 cells. The average area of clusters was 47.2 ± 6.5 μm^2^ (mean ± SEM; n = 376 clusters from 9 cells). (C) An example of formation of a cluster, which was mediated by two lysosomes undergoing directed movement (arrowheads) together with lysosomes undergoing constrained diffusion and free diffusion. The entire event was shown in Movie S6. For simplicity, only large clusters with more than 10 lysosomes were shown. Scale bar, 20 μm.

**Figure S5.**
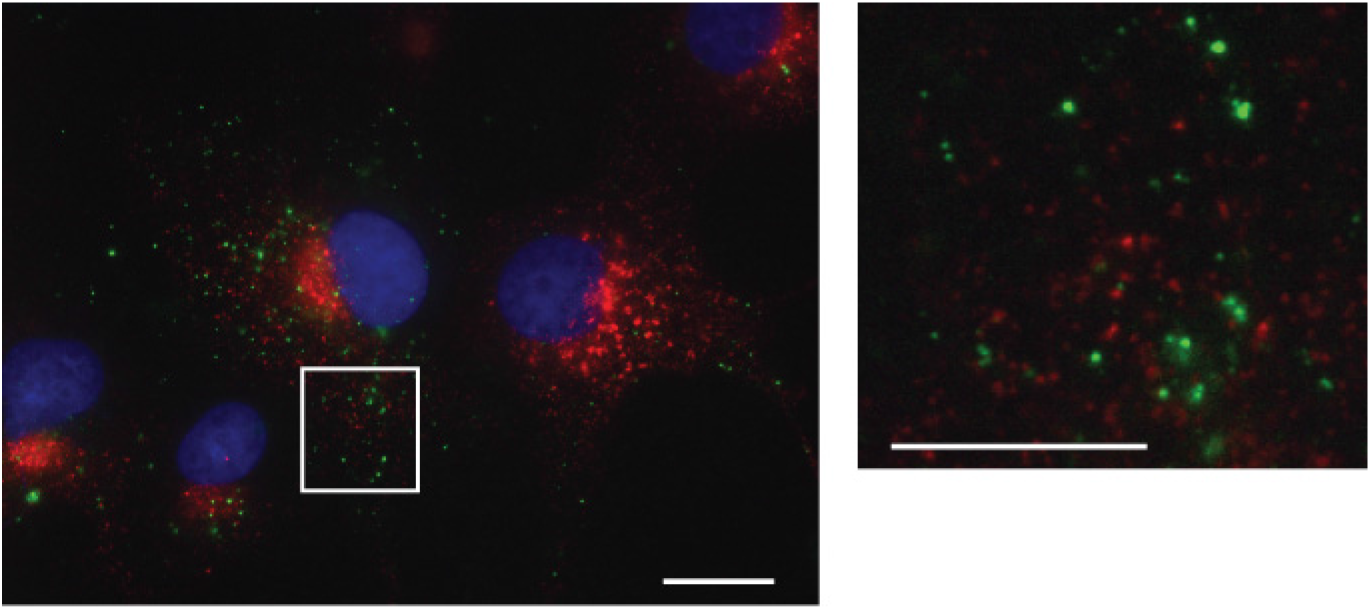
Validation of specific labeling of lysosomes as terminal endocytic compartments differentiated from late endosomes. Left panel: COS-7 cells labeled with dextran Alexa 488 for lysosome and immunostained for anti-Mannose-6-phosphate receptor (anti-M6PR). Lysosomes are defined as terminal endocytic compartments lacking M6PR (Griffiths et al., 1988; Luzio et al., 2007). They were labeled with a 3 hr pulse of dextran Alexa 488 followed by a 20 hour chase (Bright et al., 2005). Most lysosomes (green) did not colocalize with M6PR (red). Right panel: zoom-in view of the boxed area in the left panel. Scale bars: left panel, 20 μm; right panel, 10 μm.

**Figure S6.**
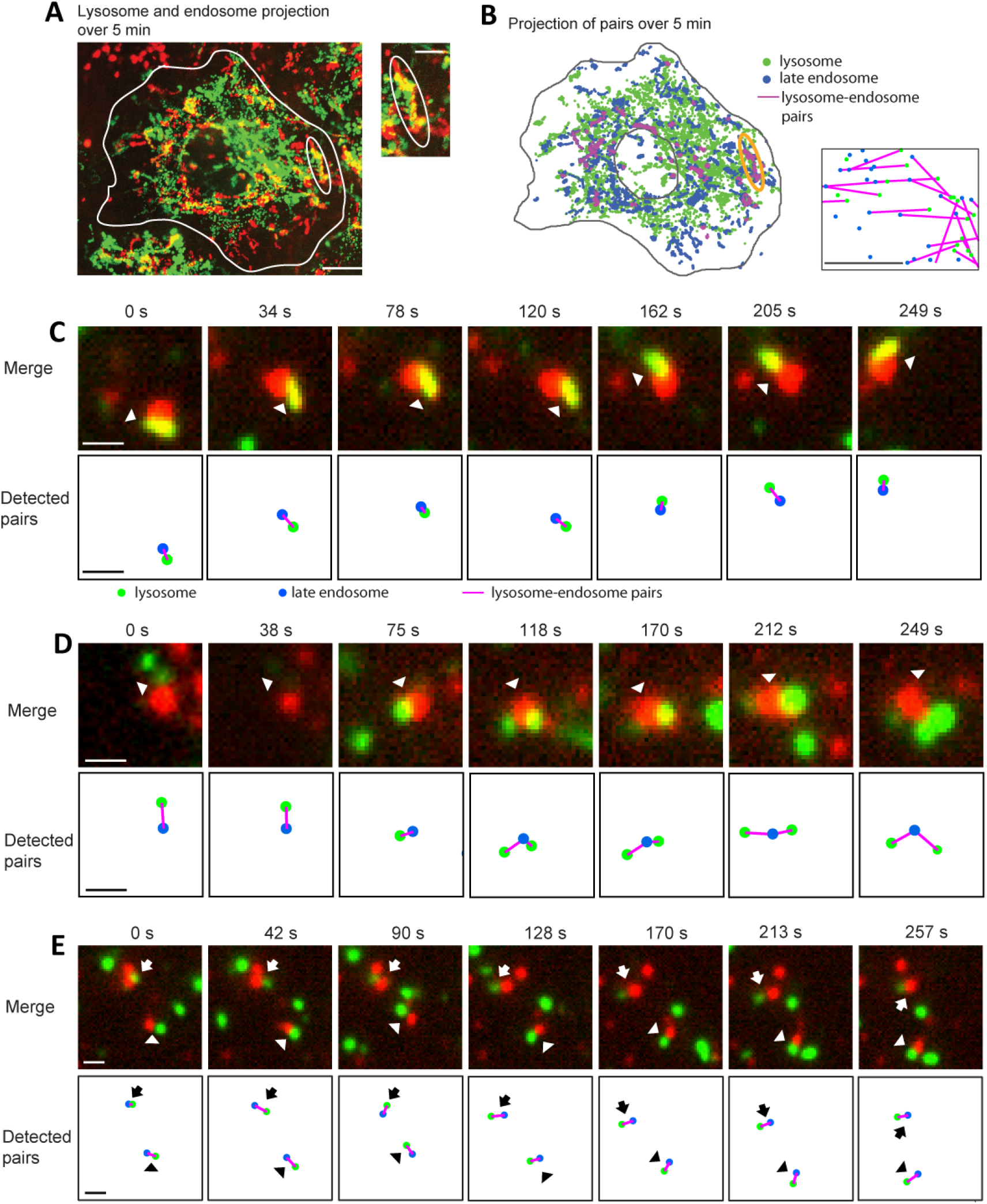
Examples of computationally detected lysosome-endosome pairs. (A) Left panel: maximum intensity projection of late endosomes (red) and lysosomes (green) of a 5-minute movie imaged at ∽5 seconds per frame. Yellow signals indicate colocalized endosome-lysosome pairs. The circled region shows the trajectory of a computationally detected endosome-lysosome pair. Inset: zoomed in view of the circled region. Scale bars: left panel, 15 μm; inset, 5 μm. (B) Left panel: maximum intensity projection of late endosomes (green) and lysosomes (blue) and detected pairs (magenta), from the same 5-minute movie as in (A). Inset: zoomed-in view of detected pairs. Scale bar: inset, 500 nm. (C-E) Various examples of computationally detected interacting lysosome-endosome pairs (lower rows) and the their actual fluorescence signals (upper rows). The pairs of organelles remained associated for at least 4 minutes. (C) An interacting lysosome-endosome pair. (D) An endosome initially interacted with one and then two lysosomes. (E) Two detected lysosome-endosome pairs close to each other. Scale bars: 1 μm.

---

**Movie S1. Extensive long-distance movement of lysosomes over the entire intracellular space.** A wide-field time-lapse movie of lysosomes in a BS-C-1 cell labeled with dextran Alexa-488 (green). Blue: cell nucleus labeled with Hoechst. The movie was taken at 4 frames per second (fps) for 1 minute. The first 20 seconds of the movie were displayed at 16 fps. Time: min:sec. Scale bar: 6.5 μm.

**Movie S2. Dynamic lysosomes in a cell patterned into a circular shape.** A wide-field time-lapse movie of lysosomes in a micro-patterned BS-C-1 cell labeled with dextran Alexa-488 (green). Blue: cell nucleus. The movie was taken at 2.5 fps for 1 minute. Every third frame was selected so that a total of 51 frames were displayed at 10 fps. Time: min:sec. Scale bar, 6.5 μm.

**Movie S3. Lysosomal dynamics before and after nocodazole treatment for depolymerization of microtubules.** A wide-field time-lapse of lysosomes in a COS-7 cell transfected with mCherry-LAMP1 (red) and imaged before and after nocodazole treatment (-5 min, 15 min, 30 min). Blue: cell nucleus. The movie was taken at 10 fps for 20 seconds and displayed at 24 fps. Time: sec. Scale bar: 10 μm.

**Movie S4. Lysosomal dynamics before and after latrunculin A treatment for depolymerization of actin filaments.** A wide-field time-lapse move of lysosomes in a COS-7 cell transfected with mCherry-LAMP1 (red) and imaged before and after latrunculin treatment (-5 min, 7.5min, 15 min). Blue: cell nucleus. The movie was taken at 10 fps for 20 seconds and displayed at 24 fps. Time: sec. Scale bar: 10 μm.

**Movie S5. Lysosomes form clusters over the entire intracellular space.** Left: a wide- field time-lapse movie of lysosomes in a COS-7 cell labeled with dextran Alexa-488 (green), taken every 25 second for one hour and displayed at 9 fps. Blue: cell nucleus. Right: color-coded spatial density plots. Black dots indicate lysosomes. Time: min:sec. Scale bar: 10 μm.

**Movie S6. Formation of lysosomal clusters mediated by lysosomes undergoing directed movement together with lysosomes undergoing constrained diffusion.** For simplicity, only large clusters with at least ten members were shown. Arrowheads point to lysosomes undergoing directed movement. Arrow points to a lysosome undergoing directed movement that contributes to merging and splitting of clusters. Time: sec. Scale bar: 20 μm. Lysosomes were tracked and classified from a widefield time-lapse movie of a COS-7 cell. The movie was taken at 10 fps for 20 seconds. A total of 24 frames were selected and displayed at 2 fps.

**Movie S7. Examples of stable lysosomal clusters with a life-time >20 seconds.** Arrows point to stable lysosomal clusters with lifetime longer than 20 seconds. Such clusters may be small, with less than ten lysosomes, and were largely composed of lysosomes undergoing constrained diffusion and free diffusion. Time: sec. Scale bar: 20 μm. The source time-lapse movie was taken at 10 fps for 20 seconds. Every fifth frame was selected. A total of 41 frames were displayed at 2 fps.

**Movie S8. Dynamic interactions between lysosomes and late endosomes.** A wide- field time-lapse movie of late endosomes labeled with dextran Alexa-594 (red) and lysosomes labeled with dextran Alexa-488 (green) in a COS-7 cell. Blue: cell nucleus. The movie was taken every 4.35 s for 5 min and displayed at 5.3 fps. Time: min:sec. Scale bar: 10 μm.

**Table S1.** Comparison of distance distributions of lysosomes before versus after nocodazole treatment;

| Relate to Fig. 3G-H |  |  |  |  |  |  |  |  |  |  |  |  |  |  |  |
| --- | --- | --- | --- | --- | --- | --- | --- | --- | --- | --- | --- | --- | --- | --- | --- |
| Cell No | Normalized inter-organelle distance distribution |  |  |  |  | Normalized to-nucleus distance distribution |  |  |  |  | Nearest neighbor distance distribution |  |  |  |  |
|  | KS test* | Median |  | RS test† p-value |  | KS test* | Median |  | RS test† p-value |  | KS test* | Median |  | RS test† p-value |  |
|  | p-value | 0 min | 30 min | Smaller | Larger | p-value | 0 min | 30 min | Smaller | Larger | p-value | 0 min | 30 min | Smaller | Larger |
| 1 | 1.6×10<sup>-4</sup> | 0.357 | 0.349 | 1.1×10<sup>-6</sup> | >0.99 | 0.026 | 0.351 | 0.432 | 0.99 | 0.013 | 0.39 | 1.80 | 1.82 | 0.54 | 0.46 |
| 2 | 1.3×10<sup>-5</sup> | 0.395 | 0.386 | 0.50 | 0.50 | 0.74 | 0.328 | 0.332 | 0.42 | 0.58 | 0.45 | 1.87 | 2.09 | 0.80 | 0.20 |
| 3 | 6.0×10<sup>-8</sup> | 0.371 | 0.353 | >0.99 | 1.4×10<sup>-6</sup> | 7.8×10<sup>-3</sup> | 0.531 | 0.353 | 6.6×10<sup>-4</sup> | >0.99 | 0.07 | 1.57 | 1.89 | 0.94 | 0.057 |
| 4 | 7.9×10<sup>-13</sup> | 0.322 | 0.313 | >0.99 | 2.31×10<sup>-65</sup> | 3.0×10<sup>-3</sup> | 0.331 | 0.270 | 2.2×10<sup>-4</sup> | >0.99 | 0.62 | 1.73 | 1.65 | 0.30 | 0.70 |
| 5 | 0.019 | 0.341 | 0.339 | >0.99 | 2.3×10<sup>-14</sup> | 0.52 | 0.397 | 0.365 | 0.18 | 0.82 | 0.73 | 1.61 | 1.53 | 0.44 | 0.56 |
| 6 | 6.0×10<sup>-8</sup> | 0.372 | 0.385 | >0.99 | >0.99 | 5.0×10<sup>-4</sup> | 0.322 | 0.427 | 0.99 | 7.8×10<sup>-3</sup> | 0.055 | 1.96 | 1.71 | 0.018 | 0.98 |
| 7 | 1.1×10<sup>-6</sup> | 0.426 | 0.413 | 3.1×10<sup>-28</sup> | >0.99 | 0.71 | 0.515 | 0.452 | 0.33 | 0.67 | 0.11 | 1.77 | 1.96 | 0.54 | 0.46 |
| 8 | 7.1×10<sup>-9</sup> | 0.325 | 0.317 | >0.99 | 8.3×10<sup>-93</sup> | 0.055 | 0.374 | 0.306 | 0.021 | 0.98 | 6.4×10<sup>-3</sup> | 1.86 | 1.56 | 0.011 | 0.99 |
| 9 | 1.4×10<sup>-4</sup> | 0.380 | 0.383 | >0.99 | 8.1×10<sup>-7</sup> | 0.92 | 0.531 | 0.495 | 0.38 | 0.62 | 0.032 | 1.68 | 1.41 | 4.9×10<sup>-3</sup> | >0.99 |
| n(p<0.05) | 9 |  |  | 2 | 6 | 5 |  |  | 3 | 2 | 2 |  |  | 3 | 0 |

\* KS test: two sample Kolmogorov-Smirnov tests to test if distance measures from two cells are from the same distribution.
† RS test: one-tailed Wilcoxon rank sum test for comparing changes in the median of the distance distributions during treatment.

**Table S2.** Comparison of distance distributions of lysosomes before versus after ciliobrevin D treatment

| Relate to Fig. 3J-K |  |  |  |  |  |  |  |  |  |  |  |  |  |  |  |
| --- | --- | --- | --- | --- | --- | --- | --- | --- | --- | --- | --- | --- | --- | --- | --- |
| Cell No | Normalized inter-organelle distance distribution |  |  |  |  | Normalized to-nucleus distance distribution |  |  |  |  | Nearest neighbor distance distribution |  |  |  |  |
|  | KS test* | Median |  | RS test† p-value |  | KS test* | Median |  | RS test† p-value |  | KS test* | Median |  | RS test† p-value |  |
|  | p-value | 0 min | 1 hour | Smaller | Larger | p-value | 0 min | 1 hour | Smaller | Larger | p-value | 0 min | 1 hour | Smaller | Larger |
| 1 | <1.0×10<sup>-299</sup> | 0.338 | 0.400 | >0.99 | 8.1×10<sup>-251</sup> | 1.4×10<sup>-4</sup> | 0.376 | 0.463 | >0.99 | 1.9×10<sup>-4</sup> | 6.7×10<sup>-3</sup> | 1.58 | 1.46 | 7.3×10<sup>-3</sup> | 0.99 |
| 2 | <1.0×10<sup>-299</sup> | 0.375 | 0.421 | >0.99 | <1.0×10<sup>-299</sup> | 7.1×10<sup>-5</sup> | 0.462 | 0.510 | >0.99 | 2.5×10<sup>-4</sup> | 2.1×10<sup>-4</sup> | 1.63 | 1.40 | 4.6×10<sup>-6</sup> | >0.99 |
| 3 | <1.0×10<sup>-299</sup> | 0.383 | 0.439 | >0.99 | <1.0×10<sup>-299</sup> | 4.3×10<sup>-6</sup> | 0.402 | 0.454 | >0.99 | 1.3×10<sup>-6</sup> | 4.9×10<sup>-4</sup> | 1.32 | 1.18 | 5.7×10<sup>-5</sup> | >0.99 |
| 4 | 1.3×10<sup>-22</sup> | 0.346 | 0.339 | 0.012 | 0.99 | 8.2×10<sup>-5</sup> | 0.415 | 0.419 | 0.99 | 0.010 | 0.17 | 1.27 | 1.20 | 0.19 | 0.81 |
| 5 | 1.3×10<sup>-45</sup> | 0.366 | 0.385 | >0.99 | 1.2×10<sup>-70</sup> | 0.025 | 0.454 | 0.502 | 0.89 | 0.11 | 7.6×10<sup>-4</sup> | 1.69 | 1.48 | 1.5×10<sup>-4</sup> | >0.99 |
| 6 | <1.0×10<sup>-299</sup> | 0.342 | 0.424 | >0.99 | <1.0×10<sup>-299</sup> | 0.12 | 0.463 | 0.482 | 0.85 | 0.15 | 8.8×10<sup>-5</sup> | 1.57 | 1.37 | 2.5×10<sup>-4</sup> | >0.99 |
| 7 | 7.4×10<sup>-64</sup> | 0.385 | 0.416 | >0.99 | 6.4×10<sup>-77</sup> | 0.014 | 0.427 | 0.470 | 0.99 | 0.011 | 0.34 | 1.76 | 1.72 | 0.22 | 0.78 |
| 8 | <1.0×10<sup>-299</sup> | 0.340 | 0.408 | >0.99 | <1.0×10<sup>-299</sup> | 1.4×10<sup>-11</sup> | 0.353 | 0.489 | >0.99 | 5.5×10<sup>-14</sup> | 9.7×10<sup>-3</sup> | 1.32 | 1.26 | 5.6×10<sup>-3</sup> | 0.99 |
| 9 | <1.0×10<sup>-299</sup> | 0.341 | 0.400 | >0.99 | <1.0×10<sup>-299</sup> | 6.0×10<sup>-5</sup> | 0.359 | 0.412 | >0.99 | 5.3×10<sup>-5</sup> | 1.5×10<sup>-4</sup> | 1.21 | 1.13 | 2.9×10<sup>-4</sup> | >0.99 |
| 10 | 2.6×10<sup>-4</sup> | 0.364 | 0.370 | >0.99 | 4.5×10<sup>-4</sup> | 5.3×10<sup>-3</sup> | 0.366 | 0.444 | >0.99 | 5.6×10<sup>-4</sup> | 0.021 | 1.50 | 1.43 | 0.021 | 0.98 |
| 11 | 2.6×10<sup>-162</sup> | 0.366 | 0.399 | >0.99 | 7.2×10<sup>-116</sup> | 0.076 | 0.442 | 0.466 | 0.96 | 0.039 | 0.25 | 1.58 | 1.63 | 0.50 | 0.50 |
| 12 | 1.0×10<sup>-90</sup> | 0.333 | 0.365 | >0.99 | 1.4×10<sup>-125</sup> | 7.7×10<sup>-3</sup> | 0.326 | 0.393 | 0.98 | 0.023 | 0.018 | 1.61 | 1.64 | 0.16 | 0.84 |
| 13 | 5.4×10<sup>-299</sup> | 0.314 | 0.351 | >0.99 | <1.0×10<sup>-299</sup> | 0.012 | 0.390 | 0.450 | >0.99 | 1.0×10<sup>-3</sup> | 0.021 | 1.56 | 1.43 | 1.9×10<sup>-3</sup> | >0.99 |
| n(p<0.05) | 13 |  |  | 1 | 12 | 11 |  |  | 0 | 11 | 10 |  |  | 9 | 0 |

\* KS test: two sample Kolmogorov-Smirnov tests to test if distance measures from two cells are from the same distribution.
† RS test: one-tailed Wilcoxon rank sum test for comparing changes in the median of the distance distributions during treatment.

**Table S3.**
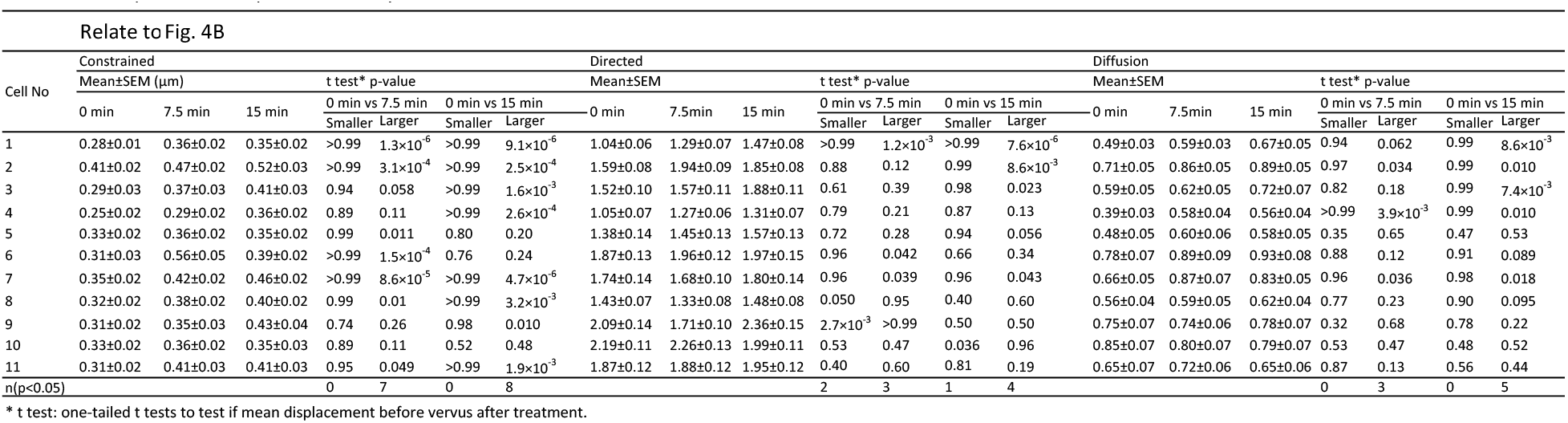
Comparison of displacement of lysosomes before versus after latrunculin A treatment

**Table S4.**
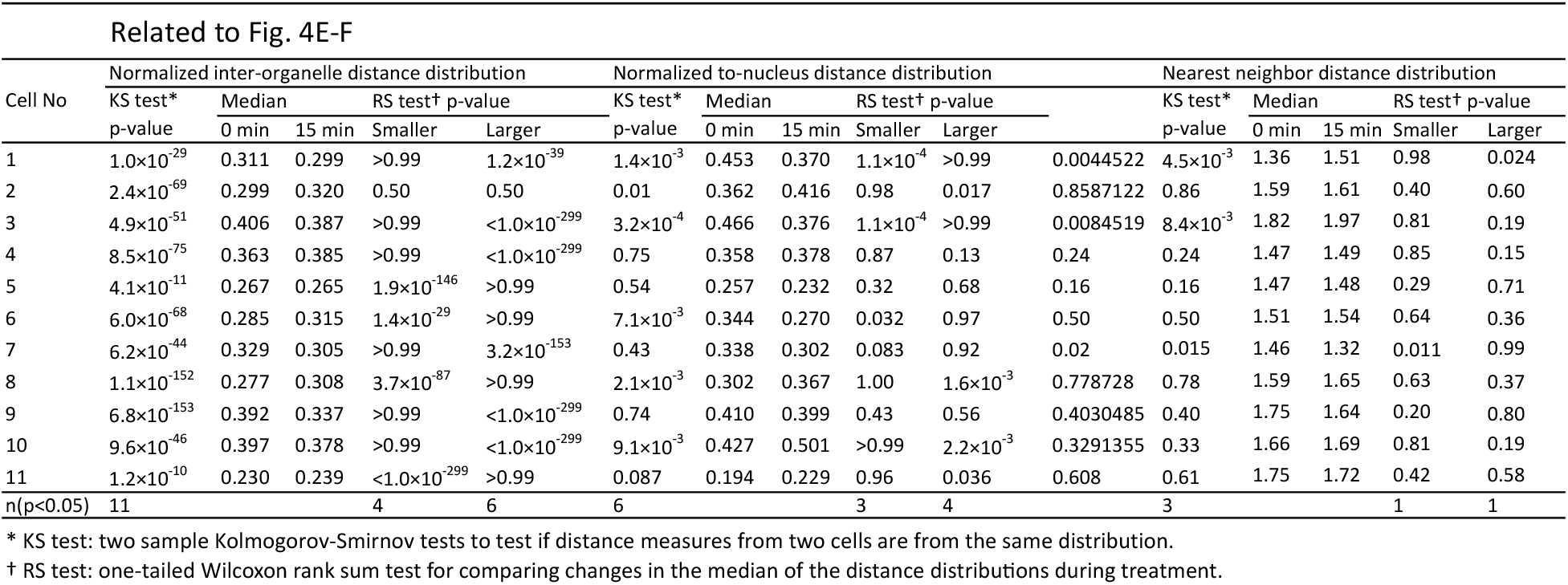
Comparison of distance distributions of lysosomes before versus after latrunculin A treatment

**Table S5.**
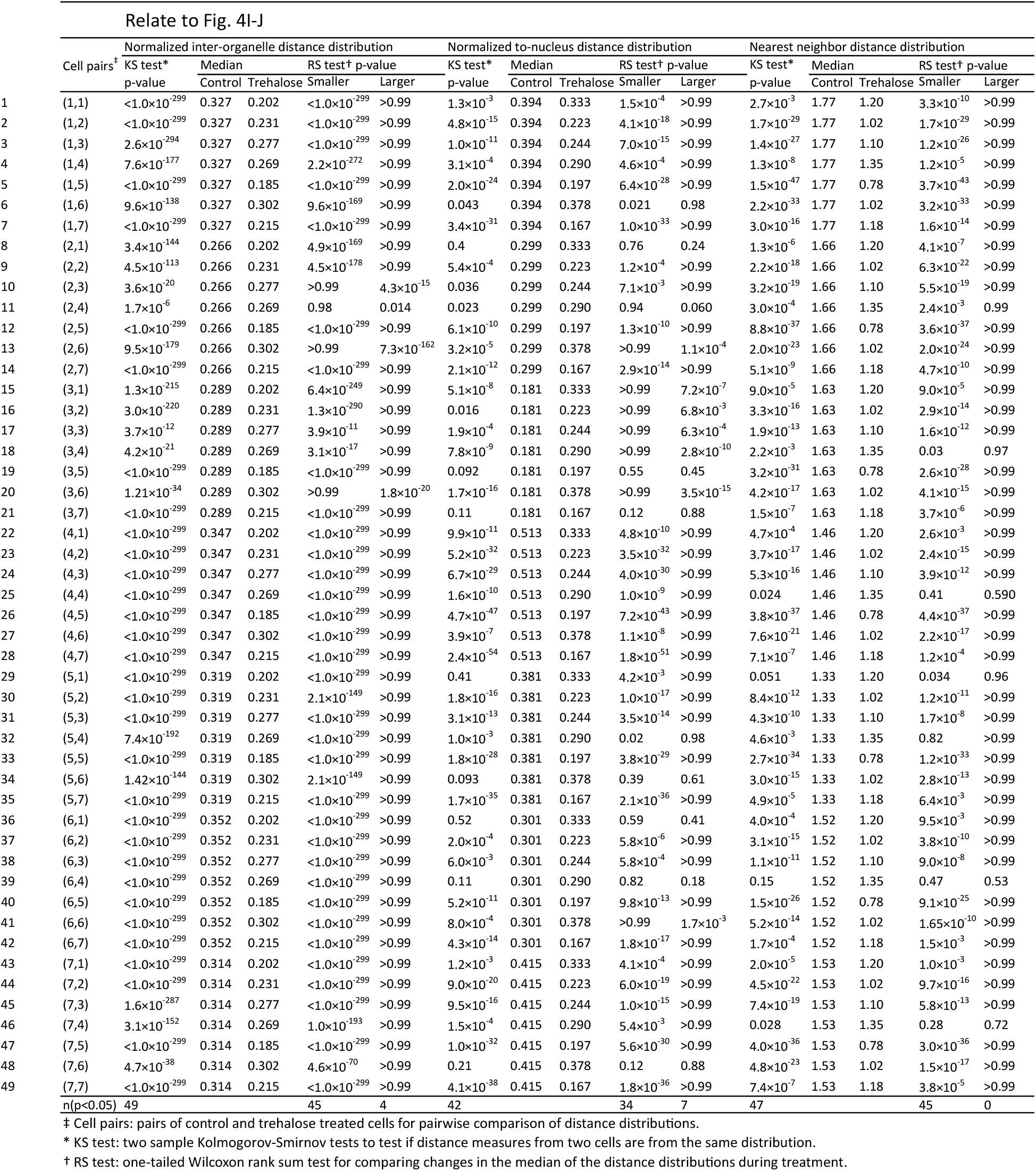
Comparison of distance distributions of lysosomes in control and trehalose treated cells

